# Cellular Determinants of CA1 Pyramidal Cell Recruitment during Sharp-Wave Ripples

**DOI:** 10.64898/2026.09.28.755157

**Authors:** Raphael Angerbauer, Tae Ho Lim, Peter Jonas, Fritjof Helmchen, Xiaomin Zhang

**Affiliations:** Brain Research Institute, University of Zurich, Zurich, Switzerland; Neuroscience Center Zurich, University of Zurich, Zurich, Switzerland; URPP Adaptive Brain Circuits in Development and Learning (AdaBD), University of Zurich, Zurich, Switzerland; Institute of Science and Technology Austria (ISTA), Klosterneuburg, Austria

**Keywords:** awake ripples, ripple diversity, hippocampus, CA1, pyramidal cells, intracellular membrane potential dynamics, *in vivo* whole-cell patch-clamp recordings

## Abstract

Hippocampal ripples, accompanied by highly synchronous neuronal firing, are critical for immediate offline memory consolidation, learning, and planning. However, the cellular mechanisms that determine which individual neurons are recruited to participate in action potential (AP) firing during ripples remain unclear. We investigated the membrane potential (V_m_) dynamics of CA1 pyramidal cells during ripples using whole-cell recordings in awake, head-fixed mice. We found that ripple participation was strongly associated with concurrent membrane-potential depolarization on an event-to-event basis, whereas between-cell differences in overall participation propensity were jointly explained by ripple-associated depolarization and intrinsic excitability. Applying this framework across CA1 cell populations revealed that differences in recruitment between superficial and deep pyramidal cells were primarily driven by ripple-associated depolarization, counteracted by excitability. Differences between place and non-place cells were similarly driven by ripple-associated depolarization, though this effect acted in the same direction as excitability. Finally, we identified the post-ripple LFP rebound as a prominent marker of ripple diversity linked to post-ripple membrane-potential hyperpolarization at the single-neuron level. Together, these findings provide a mechanistic understanding of how CA1 pyramidal cells are selectively recruited during hippocampal ripples.

## Introduction

Hippocampal sharp wave ripples (SWR) are one of the most synchronous oscillatory patterns in the mammalian brain, accompanied by highly synchronous firing of CA1 pyramidal cells. In local field potential (LFP) recordings, they are often comprised of a sharp wave component superimposed by high-frequency oscillations (80-250 Hz), known as ripples.^1^ Sharp waves result from strong excitatory inputs impinging onto the CA1 apical dendrites in the stratum radiatum (SR), whereas ripples reflect alternating peri-somatic inhibition and synchronous spikes in the stratum pyramidale (SP).^2^ A hallmark feature of ripples is the coordinated reactivation of previously encoded neural activity in a temporally compressed form during “off-line” states, such as quiet wakefulness and non-rapid eye movement (non-REM) sleep.^3,4^ Recent evidence suggests that awake replay reinstates representations of prior experiences, supporting memory consolidation, learning, and future planning.^5–10^ Disruption of awake ripples and their associated neural activity impairs performance in spatial memory tasks in rodents.^11^ Despite extensive research establishing the information content and functional role of neuronal firing during awake ripples, the precise mechanisms underlying single-neuron recruitment during awake ripples remain poorly understood.

Several pioneering studies have investigated intracellular V_m_ dynamics during ripples using sharp electrode or whole-cell patch clamp recordings in both anesthetized and awake animals.^13–16^ These studies consistently found pronounced depolarization and intracellular ripples in CA1 pyramidal cells during ripples as well as a prolonged hyperpolarization afterward. Notably, the magnitude of ripple-associated depolarization varies systematically across CA1 PCs. Superficial calbindin-positive CA1 PCs, found closer to the stratum radiatum (SR), tend to depolarize during ripples, driving greater participation, whereas deep calbindin-negative CA1 PCs, residing closer to the stratum oriens (SO), tend to hyperpolarize during ripples, reducing participation. This difference has been attributed to differential perisomatic inhibition: deep CA1 PCs receive dense inhibitory drive from parvalbumin-positive (PV+) basket cells, producing net hyperpolarization, whereas superficial CA1 PCs receive comparatively weaker local feedforward inhibitory drive, allowing excitatory input to dominate and drive net depolarization.^16^ However, CA1 PCs have been appreciated as being heterogeneous along numerous other axes, including their developmental, morphological, molecular, connectivity profiles, as well as their place coding properties. Whether intrinsic cellular properties, potentially underlying cellular heterogeneity, additionally predetermine ripple participation, beyond momentary synaptic input and local inhibition, remains unknown.

In addition to cellular diversity, hippocampal ripple diversity has been increasingly recognized.^18–21^ Recent work revealed substantial diversity in their main features, such as frequency, amplitude, duration, and coupling to sharp waves, indicating ripple waveforms span a continuum.^18,21^ This diversity is input-dependent: ripples with higher frequency and amplitude are predominantly driven by CA3 input, whereas ripples with lower frequency are mainly driven by input from the entorhinal cortex layer 3 (EC3) and more likely to form long ripples and ripple bursts.^19,21,22^ Very recent work revealed that ripple subtypes that occur during sleep also differ in mnemonic functions: CA3 input–driven (Rad^sink^) ripples preferentially reactivate motifs of the immediately preceding exploration (recent) experiences by recruiting both superficial and deep CA1 PCs, while EC input-driven (LM^sink^) ripples primarily reactivate motifs that predate exploration (remote) experiences by engaging deep CA1 cells.^19^ Moreover, CA3 input-driven ripples (classical high frequency ripples) are associated with neocortical metabolic up-regulation, whereas EC input-driven ripples (non-classical low frequency ripples) exhibit the opposite pattern in anesthetized rhesus monkeys.^20^ These findings suggest ripple diversity during sleep may coordinate memory reactivation within the hippocampus and direct information transfer to cortical regions for long-term storage.^20^ However, it remains unknown whether similar ripple diversity is also present during quiescent wakeful state, and how distinct awake ripple subtypes selectively recruit CA1 PCs. Resolving this question is critical for understanding how neuronal replay sequences are generated across ripple subtypes and how ripple diversity shapes memory processing.

By combining whole-cell patch-clamp with extracellular LFP recordings, we simultaneously monitored the sub- and suprathreshold activity of CA1 pyramidal cells during hippocampal ripples in awake, behaving mice. We show that differences in ripple participation between superficial and deep CA1 PCs, and between place and non-place cells, depend on an interplay between ripple-associated membrane-potential dynamics and intrinsic excitability. We further identify post-ripple LFP rebound as a prominent feature of ripple diversity linked to hyperpolarization on a single-cell level. These mechanistic insights are crucial for understanding how different subtypes of CA1 pyramidal cells are differentially recruited for neuronal replay across awake ripples, supporting learning, planning, and memory consolidation.

## Results

### Ripple-participation is associated with concurrent V_m_-depolarization in an event-to-event basis

To investigate how CA1 pyramidal cells are recruited during SWRs, we simultaneously measured the membrane potential (V_m_) of CA1 pyramidal cells (n=48) and the local field potential (LFP) using whole-cell patch-clamp and extracellular recordings in awake, head-fixed and behaving mice (Figure 1A). The patch pipette was filled with biocytin, which enabled post-hoc morphological reconstruction of a subset of the measured cells (Figure 1A, top; n = 16). Ripple events (n = 4205) were identified in the LFP trace and manually validated (Figure 1B, top, shaded regions). We observed a median ripple amplitude of 0.18 mV (IQR: [0.13, 0.24]), a median duration of 39 ms (IQR: [30, 53]), and a median frequency of 130 Hz (IQR: [123, 139]), consistent with previous reports (Figure 1C). We further observed a post-ripple LFP positive deflection (Rebound) lasting up to 200 ms following ripple offset, with a median amplitude of 78.4 μV (IQR: [64.2, 91.8]) (Figure 1D; Figure 1E; Figure S1A).

**Figure 1:**
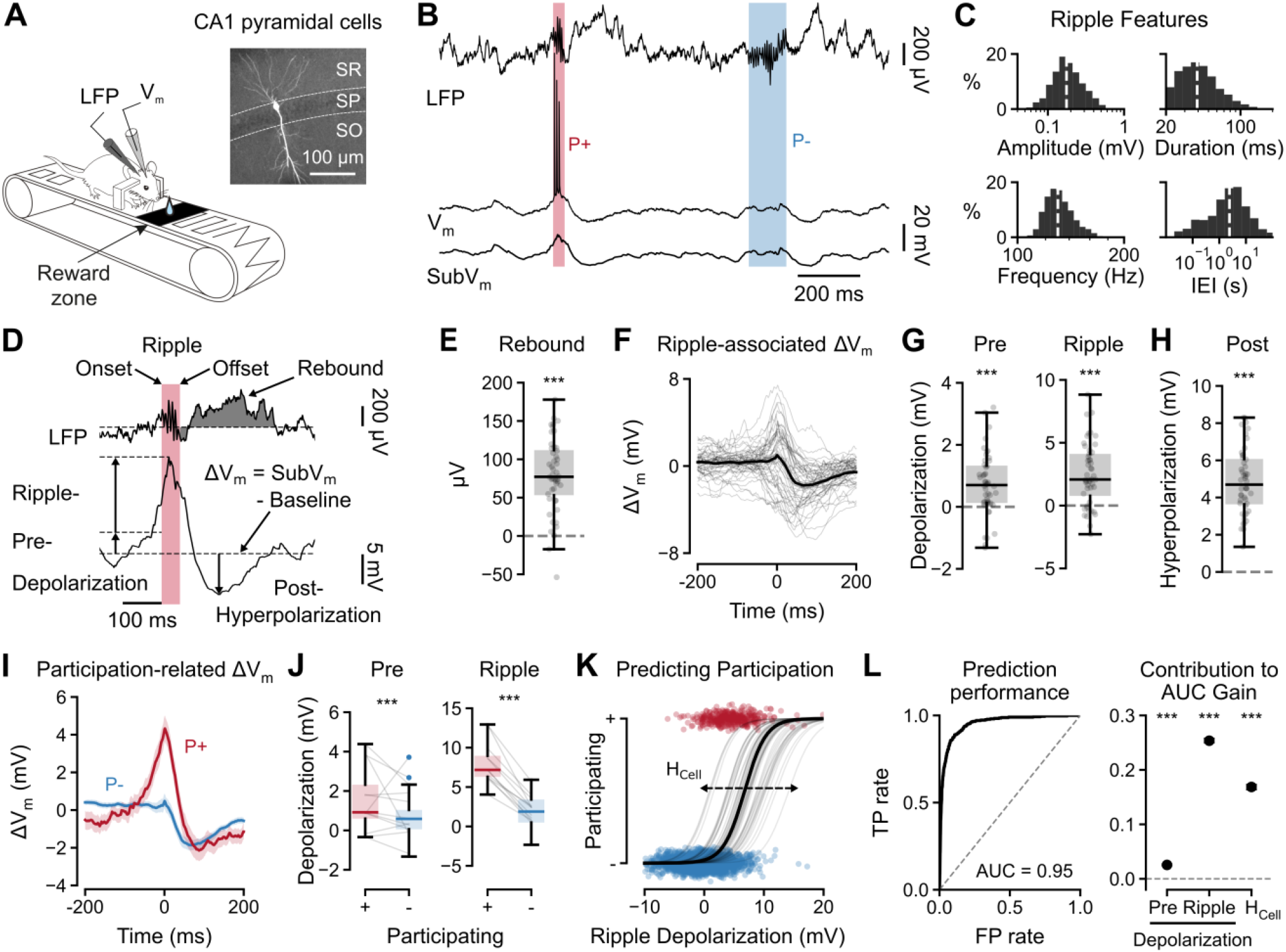
Membrane-potential dynamics underlying CA1 pyramidal-cell participation during hippocampal ripples. **(A)** Experimental configuration for intracellular recording of CA1 pyramidal cells in head-fixed mice running on a linear treadmill. Inset, representative recorded CA1 pyramidal cell, stained with biocytin, and hippocampal sublayers (SR, stratum radiatum; SP, stratum pyramidale; SO, stratum oriens). **(B)** Representative simultaneous LFP (top), membrane potential (V_m_) and subthreshold membrane-potential (SubV_m_, with APs removed) recordings. Example ripples in which the neuron participated (P+, red) or did not participate (P-, blue) are highlighted. **(C)** Distributions of ripple amplitude, duration, frequency, and inter-event interval (IEI) across detected ripple events, shown on logarithmic axes. Median is indicated as a white dashed line. **(D)** Definition of ripple-related variables. Ripple onset and offset were determined from the LFP (ripple period highlighted in red). Post-ripple rebound and the window for its analysis (ripple offset to 200-ms thereafter) are highlighted. Baseline membrane potential (median of SubV_m_ from5 s preceding onset to 5 s following offset, excluding ripple periods) was subtracted from SubV_m_ to yield the ripple-evoked depolarization (ΔV_m_). From this signal, 3 quantities were measured: pre-ripple depolarization (Pre Depolarization: mean ΔV_m_ between 50 ms and 5 ms before ripple onset), intra-ripple depolarization (Ripple Depolarization: maximum ΔV_m_ during the ripple window minus Pre-depolarization), and the post-ripple hyperpolarization (Post-hyperpolarization; Baseline minus the minimum V_m_ between 5 ms to 100 ms following ripple offset). **(E)** Distribution of post-ripple rebound amplitudes. Rebound amplitude was defined as the mean LFP in the window highlighted in D. **(F)** Ripple-triggered ΔV_m_ traces aligned to the ripple peak. Thin lines indicate individual cells, and the black line indicates the population average. **(G)** Cell-wise pre-ripple and ripple depolarization illustrating depolarization that builds before and peaks during ripples. **(H)** Cell-wise post-ripple hyperpolarization amplitude, showing that ripples are followed by hyperpolarization. **(I)** Ripple-associated ΔV_m_ (mean±SEM) for participating (P+, red) and non-participating (P-, blue) events averaged across cells. **(J)** Comparison of pre-ripple depolarization and ripple depolarization between P+ and P-events across cells. **(K)** Relationship between Ripple Depolarization and the probability of participation. Points show individual ripple events and curves show cell-specific participation functions estimated using hierarchical logistic regression; horizontal variations among curves reflect cellular heterogeneity in participation propensity (H_cell_). **(L)** Left, receiver-operating-characteristic (ROC) curve for prediction of ripple participation (area under the curve, AUC = 0.95; chance level = 0.5). Right, contribution of Pre- and Ripple Depolarization, and cell-to-cell differences in participation (H_cell_) to predictive performance. Statistical evidence is denoted by asterisks according to the highest credible level whose highest density interval excludes the null (*, 95%; **, 99%; ***, 99.9%; ns, otherwise).

We hypothesized that ripple-associated subthreshold V_m_ dynamics might influence whether a cell participates during a given ripple. To isolate the subthreshold component (SubV_m_), we removed the AP waveforms by linear interpolation from the V_m_ trace (Figure 1B, bottom) and referenced the resulting signal relative to a local baseline, which was defined as the median SubV_m_ in a symmetric 10-s window around each ripple (ΔV_m_; Figure 1D; see Methods). Ripple-peak-aligned ΔV_m_ varied substantially across cells (Figure 1F) but revealed a common temporal structure: On average, cells were slightly depolarized before ripple onset (Pre-depolarization; 0.47 mV, 95% highest-density interval (HDI): [0.24, 0.71]; Figure 1D, left; Figure 1G, left), more strongly depolarized during the ripple (Ripple-depolarization; 2.73 mV, HDI: [1.82, 3.18]; Figure 1G, right), and strongly hyperpolarized following ripple offset (Post-hyperpolarization; 4.48 mV, HDI: [3.97, 4.93]; Figure 1H), consistent with previous findings.^13,14^

To determine how SubV_m_ dynamics are related to ripple recruitment, we classified each ripple as participating (P+) if at least one AP was evoked during the ripple window, and non-participating (P-) otherwise (Figure 1B, centre). We found that P+ events were associated with significantly greater depolarization compared to P-events for pre-depolarization (0.74 mV difference, HDI: [0.44, 1.04]) and ripple-depolarization (Δ: 5.42 mV, HDI: [5.1, 5.72]; Figure1I, J) with no difference for Post-hyperpolarization (Δ: 0.06 mV, HDI: [−0.34, 0.46]; Figure1I, Figure S1B). We next asked whether pre- and ripple-depolarization were sufficient to predict participation on a ripple-by-ripple basis within individual cells. We therefore fitted a hierarchical logistic regression model predicting participation from pre- and ripple-depolarization, with a random intercept per cell to capture baseline differences in participation propensity. We found that while both pre- (β: 0.58, HDI: [0.43, 0.72]) and ripple-depolarization (β: 2.20, HDI: [1.99, 2.40]) held significant predictive power (Figure 1K and Figure S1C, D), the model also revealed substantial cell-to-cell differences in participation propensity (H_cell_; Figure 1K). To quantify the relative importance of these components, we assessed model performance using the receiver-operating-characteristic (ROC) area-under-the-curve metric (AUC: 0.948, HDI: [0.944, 0.952]) and decomposed it into the improvements in AUC attributable to pre-depolarization, ripple-depolarization, and the cell-specific component using a Shapley decomposition (Figure 1L; see Methods). Ripple-depolarization contributed the largest improvement in predictive performance (ΔAUC: 0.255, HDI: [0.25, 0.257]), followed by H_cell_ (ΔAUC: 0.169, HDI: [0.164, 0.172]), whereas pre-depolarization made only a small additional contribution (ΔAUC: 0.025, HDI: [0.025, 0.026]). Thus, while ripple-associated depolarization explained a substantial part of ripple-by-ripple variation in participation within cells, clear differences in participation propensity remained between individual cells.

### Ripple-participation propensity is jointly determined by ripple-associated depolarization and intrinsic excitability

To investigate why cells differed in their participation propensity during ripples, we calculated each cell’s participation ratio (PR), defined as the percentage of ripples in which the cell participated. PR varied substantially across cells (Figure 2A). Given the strong relationship between ripple-associated depolarization and participation on an event-to-event basis within cells, we hypothesized that differences in depolarization might similarly contribute to differences in PR between cells. However, mean ripple-associated ΔV_m_ waveforms did not show a clear, monotonic relationship with PR across quintiles (Figure 2B). To quantify how much of the cell-to-cell variation in PR could be explained by differences in depolarization, we fit a binomial regression model predicting each cell’s PR from its mean pre- and ripple-depolarization. Larger ripple-depolarization was associated with higher PR (β: 0.36, HDI: [0.23, 0.48]; Figure 2C, left) whereas stronger pre-ripple depolarization was predicted to lead to lower participation (β: −0.108, HDI: [−0.208, −0.002]; Figure 2C, right). However, only a small fraction of the cell-to-cell variance in PR was explained by pre- and ripple-depolarization (7.08%, HDI: [2.74, 12.90]; Figure 2D).

**Figure 2:**
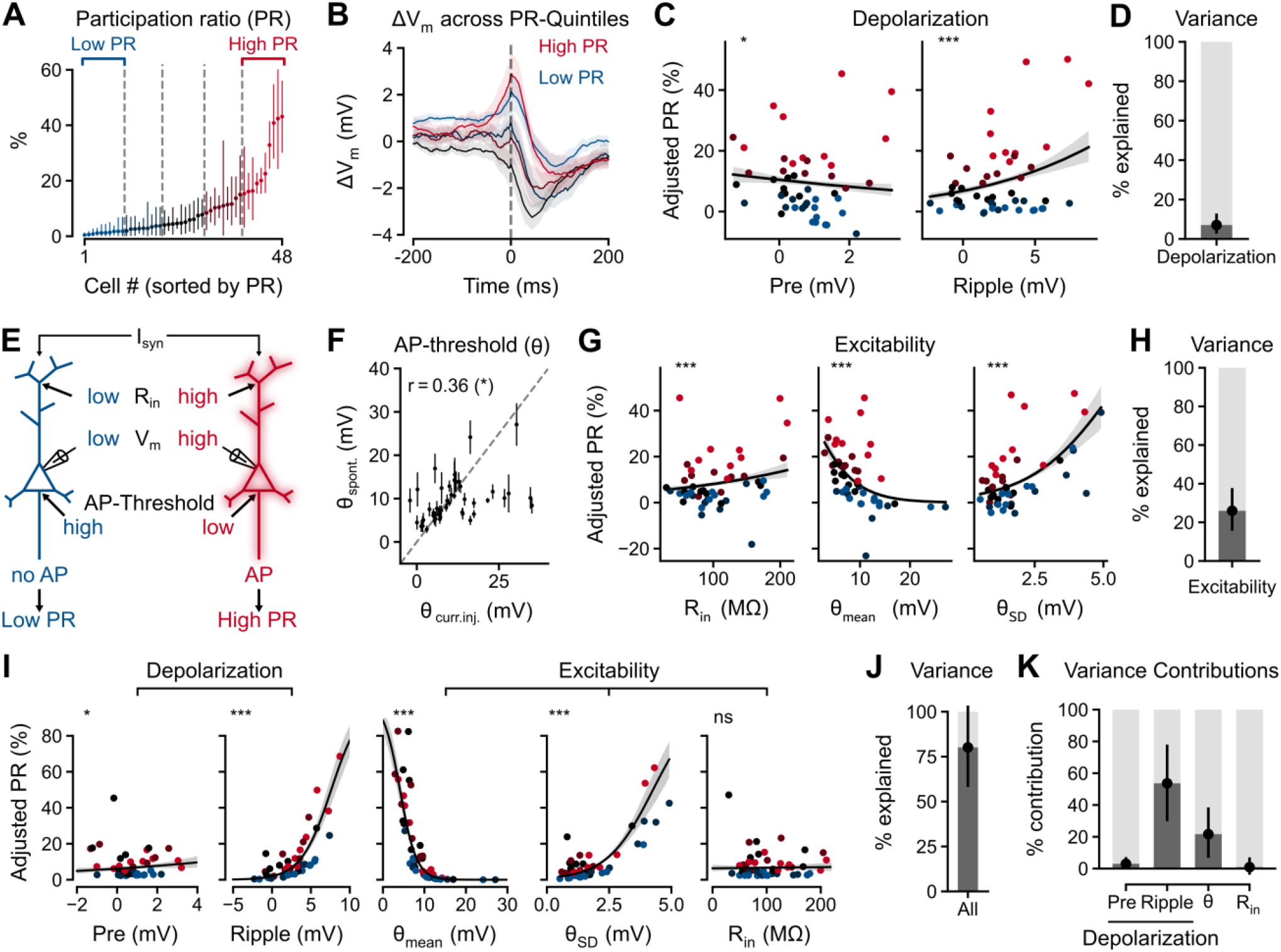
Ripple-associated depolarization and intrinsic excitability explaining cell-to-cell variability in ripple participation. **(A)** Participation ratio (PR; Median, 95% highest density interval) across CA1 pyramidal cells, estimated using hierarchical binomial regression. Cells were sorted from lowest to highest PR. Cells were split into quintiles (blue to red, separated by dashed lines) with the lowest and highest PR groups defined as Low PR and High PR cells. **(B)** Ripple-triggered membrane-potential change (ΔV_m_, mean±SE) aggregated first within cells and then within PR-quintiles. Low PR (blue) and high PR (red) cells are highlighted. **(C)** Prediction of PR using depolarization variables in a binomial regression model. Scattered points represent each cell’s PR corrected for the effect of all other predictors and are colored by PR-quintile membership. Black line represents median posterior prediction with the 95% highest density interval highlighted in grey. Significance is indicated at the top. **(D)** Fraction of the total cell-to-cell variance in PR explained by the depolarization model in C. Point and error bar indicate posterior median and 95% highest density interval. **(E)** Schematic illustration of candidate cellular determinants of ripple participation. Two cells with similar synaptic input (I_syn_) may participate differently due to differences in input resistance (R_in_) leading to increased membrane potential deflections (V_m_), or due to different AP thresholds. **(F)** Comparison of the AP-threshold estimated from spontaneous activity (θ_spont_) to the threshold measured during current injections at the start of the recording (θ_curr.inj._). Error bars indicate the standard deviation of the threshold distribution (θ_SD_) whereas the dots indicate its mean (θ_mean_). The majority of cells lying on the diagonal (dashed line) indicate that θ_spont_ agrees with θ_curr.inj._in all but a few cases. θ_spont_ is used for the rest of the study. **(G)** Prediction of PR using excitability variables θ_mean_, θ_SD_, and R_in_ in a binomial regression model. Scattered points represent each cell’s PR corrected for the effect of all other predictors and are colored by PR-quintile membership. Black line represents median posterior prediction with the 95% highest density interval highlighted in grey. Significance of the fitted coefficient is indicated at the top. **(H)** Fraction of the total cell-to-cell variance in PR explained by the excitability model in G. Point and error bar indicate posterior median and 95% highest density interval. **(I)** Prediction of PR using both depolarization and excitability variables in a binomial regression model. Scattered points represent each cell’s PR corrected for the effect of all other predictors and are colored by PR-quintile membership. Black line represents median posterior prediction with the highest density interval highlighted in grey. Significance of the fitted coefficient is indicated at the top. **(J)** Fraction of the total cell-to-cell variance in PR explained by the combined depolarization and excitability model in I. Point and error bar indicate posterior median and 95% highest density interval. **(K)** Shapley decomposition of the cell-to-cell variance in PR explained by the full model, showing the contribution attributable to each predictor. Points and error bars indicate posterior median and 95% highest density interval. Statistical evidence is denoted by asterisks according to the highest credible level whose highest density interval excludes the null (*, 95%; **, 99%; ***, 99.9%; ns, otherwise).

These results suggest that cell-to-cell differences in participation are not fully explained by depolarization alone, suggesting that differences in intrinsic excitability might additionally contribute. We therefore hypothesize that two cells receiving similar synaptic input (I_syn_) may participate differently due to differences in intrinsic excitability: higher input resistance (R_in_) would produce larger membrane potential (V_m_) deflections for the same input, and a lower AP threshold would make that deflection more likely to cross threshold. Therefore, R_in_ and AP-threshold would jointly determine whether a given synaptic input is converted into an AP and thus into ripple participation (Figure 2E). We measured each cell’s R_in_ and AP threshold, defined as membrane potential at AP onset relative to resting membrane potentials preceding current injection, using a series of hyperpolarizing and depolarizing current injections at the start of the recording (see Methods). However, in a small subset of cells the AP threshold obtained from current injection were unusually high and did not correspond well to AP threshold measured from baseline-corrected spontaneous activity (Figure S2A). This is likely due to access resistance fluctuations during running cannot be perfectly compensated by bridge balance in real time in head-fixed running mice, adding variability to current-injection AP threshold estimates that is unrelated to excitability. We therefore attempted to estimate the AP threshold from spontaneous, baseline-corrected activity instead. As the ΔV_m_ at-onset varied quite substantially between APs within each cell, we modelled θ to be normally distributed, parametrized by a cell-specific mean (θ_mean_) and standard deviation (θ_SD_; Figure S2B). We estimated both θ_mean_ and θ_SD_ for each cell using a hierarchical Bayesian Poisson model that additionally accounted for spike-history effects (Figure S2B–D; see Methods). As an independent validation of these estimates, we compared the AP threshold estimated from spontaneous activity with the directly measured AP threshold obtained during current injection. (Figure 2F). Both values were significantly correlated (r = 0.36, p = 0.01) and agreed in most cases, except for those aforementioned cells that showed unusually high thresholds during current injection.

We next asked whether these intrinsic excitability measures could account for cell-to-cell differences in participation, by fitting a binomial regression model using R_in_, θ_mean_ and θ_SD_ as predictors. As hypothesized, R_in_ showed a positive association with PR (β: 0.26, HDI: [0.15, 0.38]; Figure 2G, left and centre), whereas the relationship between estimated θ_mean_ and PR was negative (β: −0.93, HDI: [−1.13, −0.73]; Figure 2G, centre). In addition, cells with high θ_SD_ were on average associated with higher PR (β: 0.74, HDI: [0.59, 0.89]; Figure 2G, right), indicating that in these cells the threshold is more often sufficiently low for the cell to spike. Furthermore, all three variables combined accounted for a significant fraction of the between-cell variance (25.70%, HDI: [15.73, 37.38]; Figure 2H). Nevertheless, neither depolarization nor any individual measure of intrinsic excitability was sufficient to account for the full range of participation across cells, suggesting that cell-to-cell differences in PR arise from the combined influence of depolarization and excitability factors.

We therefore combined the depolarization and intrinsic excitability measures in a single binomial regression model to determine their joint contribution to between-cell variation in PR. After accounting for the other predictors in the model, pre-depolarization, ripple-depolarization, θ_mean_, and θ_SD_ remained significantly associated with PR, whereas R_in_ showed no significant association, suggesting that the effect of R_in_ on participation was indeed mediated through V_m_ changes (Figure 2I, right). Accounting for all other variables, pre-ripple depolarization had a positive association with PR instead of negative as before (Figure 2I, left). Together, the full set of predictors accounted for a high fraction of the cell-to-cell variance in participation (80.6%, HDI: [59.1, 104.4]; Figure 2J). Decomposing the variance attributable to the individual factors revealed that differences in ripple-associated depolarization accounted for the largest contribution (53.7%, HDI: [29.8, 78.6]), followed by AP threshold (22%, HDI: [6.7, 38.6]), whereas pre-ripple depolarization and R_in_ contributed comparatively little (Figure 2K). Thus, cell-to-cell differences in ripple participation were best accounted for by the combined effects of ripple-associated depolarization and intrinsic excitability, rather than by any single cellular property alone.

### Ripple-associated depolarization outweighs opposing threshold effects to increase participation of superficial CA1 pyramidal cells

Having identified depolarization and intrinsic excitability as joint determinants of cell-to-cell differences in participation, we next asked whether these mechanisms also correlate with differences between established CA1 pyramidal cell subclasses, specially, superficial and deep cells, and place and non-place cells. Superficial cells (SC) are located closer to SR, whereas deep cells (DC) locate closer to SO (Figure 3A). DCs have previously been reported to have higher firing rate, higher burstiness, lower resting membrane potential (RMP) and distinct theta phase-locking properties compared to SCs.^23^ Importantly, SC have also been shown to participate more often during ripples than DC^16,24^, but what specific cellular properties drive this difference remains unclear. To address this question, we classified the biocytin-filled cells (n = 16; Figure 3A) as SC and DC according to their relative somatic position in the CA1 pyramidal layer (Figure S3A). Consistent with previous results, SC had a lower average firing rate (rate-ratio: 0.22, HDI: [0.04, 0.54]; Figure S3B) and a lower likelihood of bursting (odds-ratio: 0.11, HDI: [0.06, 0.16]; Figure S3C), however both RMP (Δ: 1.09 mV, HDI: [−5.79, 6.90]; Figure S3D) and preferred theta phase (Δ: 14.14°, p: 0.9) did not differ between the cell types (Figure S3E). Furthermore, SC showed higher ripple participation than DC (ΔPR: 8.8 %; odds-ratio: 1.9, HDI: [1.2, 2.8]; Figure 3B), although it accounted for only a small fraction of the overall cell-to-cell variation in PR (3.1 %, HDI: [0, 7.8]; Figure 3C).

**Figure 3:**
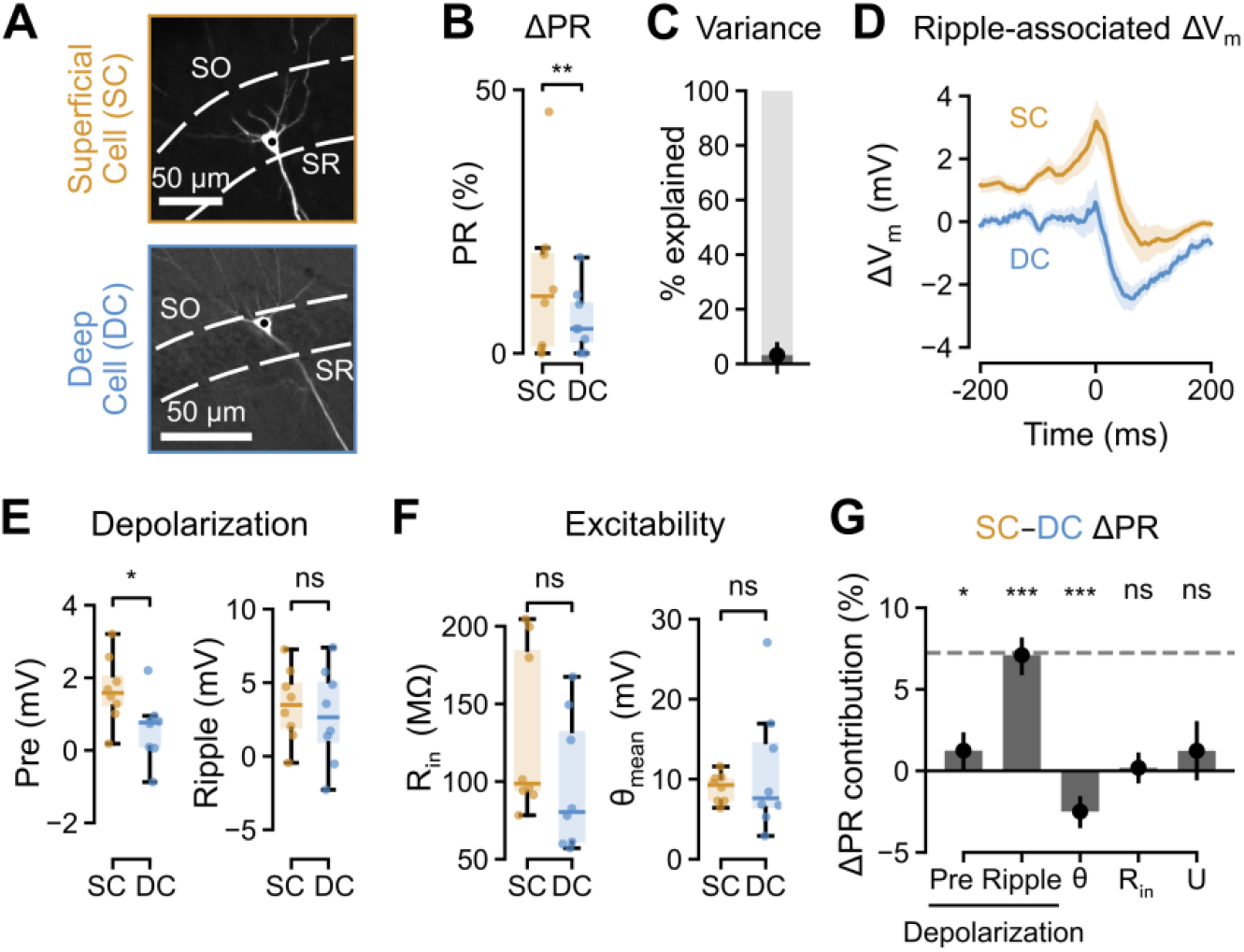
Ripple-associated depolarization and intrinsic excitability competing to explain increased participation during ripples in superficial CA1 pyramidal cells. **(A)** Representative superficial (SC, orange) and deep (DC, blue) CA1 pyramidal cells of those filled with biocytin during the recording (n = 16). Dashed lines indicate the pyramidal-cell layer boundary; SO, stratum oriens; SR, stratum radiatum. **(B)** Participation ratio difference (ΔPR) between superficial and deep cells, tested using binomial regression. Note that superficial cells showed higher ripple participation than deep cells. **(C)** Fraction of between-cell variance in PR explained by superficial–deep identity. Points and error bars indicate posterior median and 95% highest density interval. **(D)** Ripple-associated membrane-potential change (ΔV_m_, mean±SE) aggregated first within cells and then within cell types. **(E)** Difference of depolarization variables between cell types, tested using robust Student-T regression. Note that superficial cells show V_m_ depolarization building up before ripple onset. **(F)** Difference of excitability variables between superficial and deep cells, tested using robust Student-T regression. Note that excitability variables show no significant differences between cell types. **(G)** Shapley decomposition of the difference in PR between superficial and deep cells explained by the full model, showing the contribution attributable to each predictor. The fraction unexplained by the model (U) is shown on the left. Points and error bars indicate posterior median and 95% highest density interval. Statistical evidence is denoted by asterisks according to the highest credible level whose highest density interval excludes the null (*, 95%; **, 99%; ***, 99.9%; ns, otherwise).

We next asked whether the higher participation of SC could be explained by differences in ripple-associated depolarization. Ripple-associated ΔV_m_ responses differed between the two groups, with SCs showing greater pre-ripple depolarization than DCs (Δ: 1.05 mV, HDI: [0.14, 1.99]), whereas ripple-depolarization was similar between SCs and DCs (Δ: 0.72 mV, HDI: [−2.39, 3.68]; Figure 3D, E). Investigating how intrinsic excitability might differ between the cell types, neither R_in_ nor θ_mean_ differed significantly between SCs and DCs (Δ: 33.3 MΩ, HDI: [−22.2, 83.9]; Δ: −2.02 mV, HDI: [−8.22, 3.87]; Figure 3F). However, as demonstrated above, differences between cells arise not from individual factors alone but due to a combination of depolarization and intrinsic excitability. To gauge the relative importance of each factor in contributing to participation difference between SCs and DCs, we iteratively added predictors to the combined depolarization-excitability model fitted previously (Figure 2I) and calculated the ΔPR using the model’s predictions. Averaging the individual contributions of each variable (Figure 3G; see Methods, Shapley Decomposition) we found that ripple-depolarization was predominantly responsible for the observed participation differences between SCs and DCs (ΔPR contribution: 7.1 %, HDI: [5.9, 8.2]), with pre-depolarization contributing to this difference as well (ΔPR contribution: 1.2 %, HDI: [0.1, 2.4]). AP threshold θ showed a negative contribution to the observed ΔPR (ΔPR contribution: −2.5 %, HDI: [−3.5, −1.6], whereas R_in_ and the unexplained residual (U) contributed very little to the observed differences (ΔPR contribution: 0.2 %, HDI: [−0.8, 1.2]; ΔPR contribution: 1.2 %, HDI: [−0.6, 3.1]). These results indicate that although the SC’s AP-threshold would make them less likely to participate than DC at the same level of depolarization, the stronger depolarization of SC during ripples compensates for this disadvantage, resulting in a higher participation propensity.

### Place cells show lower ripple participation due to reduced ripple-associated depolarization

Having examined how ripple participation relates to an anatomical classification of CA1 pyramidal cells, we next asked whether similar principles apply to functionally defined populations, specifically place cells vs. non-place cells. During spatial navigation, a subset of CA1 pyramidal cells forms place fields and increases its firing when the animal occupies specific locations in the environment. These place cells (PCs) are often later reactivated during ripples in replay events.^3,25^ However, whether PCs differ systematically from non-place cells (NPCs) in their overall propensity to participate in ripples and, if so, which cellular mechanisms underlie this difference, remains unclear. We therefore classified cells as PCs or NPCs based on their spatial firing during locomotion (Figure 4A; Figure S4A; see Methods). The detected PCs showed similar firing rates (rate-ratio: 1.13, HDI: [0.46, 1.94]; Figure S4B) and RMP (Δ: 0.78 mV, HDI: [−4.90,5.51]; Figure S4D) compared to NPCs but were more likely to burst (odds-ratio: 2.44, HDI: [2.19, 2.73]; Figure S4C). The two cell types also did not significantly differ in their propensity to phase-lock to the hippocampal theta rhythm (odds-ratio: 1.71, p = 1.0; Figure S4E, left) nor in their preferred phase in the theta cycle (Δ: 108.11°, p: 0.27; Figure S4E, right). Comparing their ripple participation propensity, we found that PCs participated significantly less during ripples than NPCs (ΔPR: −12.7 %; odds-ratio: 0.33, HDI: [0.24, 0.42]; Figure 4B), a difference that accounted for 26.8 % (HDI: [12.9, 42.8]) of the total cell-to-cell variability (Figure 4C).

**Figure 4:**
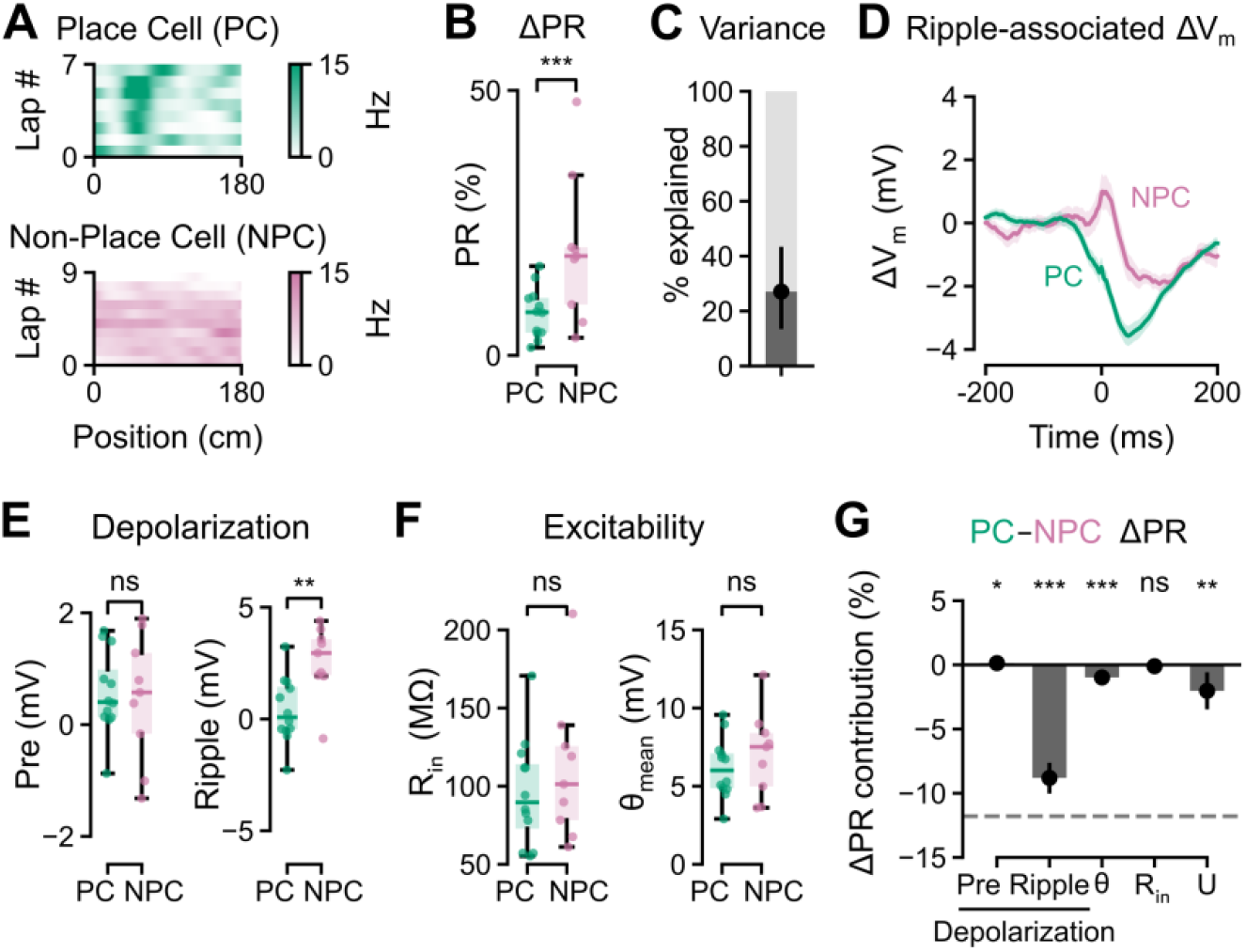
Reduced ripple-associated depolarization leading to lower participation during ripples in place cells. **(A)** Representative spatial firing-rate maps across individual laps on the linear track of an example place cell (PC) and an example non-place cell (NPC). Color indicates firing rate (Hz). **(B)** Difference in participation ratio (ΔPR) between place cells and non-place cells, tested using binomial regression. Note that place cells showed significant lower PR than non-place cells. **(C)** Fraction of between-cell variance in PR explained by place–non-place cell classification. Points and error bars indicate posterior median and 95% highest-density interval. **(D)** Ripple-associated membrane-potential change (ΔV_m_, mean±SE) aggregated first within cells and then within cell types. **(E)** Difference in depolarization variables between place cells and non-place cells, tested using robust Student-T regression. Note that non-place cells showed significantly greater depolarization during ripples. **(F)** Difference in excitability variables between place cells and non-place cells, tested using robust Student-T regression. Note that excitability variables show no significant differences between cell types. **(G)** Shapley decomposition of the difference in PR between place and non-place cells explained by the full model, showing the contribution attributable to each predictor. The fraction unexplained by the model (U) is shown on the left. Points and error bars indicate posterior median and 95% highest density interval. Statistical evidence is denoted by asterisks according to the highest credible level whose highest density interval excludes the null (*, 95%; **, 99%; ***, 99.9%; ns, otherwise).

We next asked whether the lower participation of PCs could be explained by differences in ripple-associated depolarization or intrinsic excitability and found that ripple-associated ΔV_m_ responses differed between the two groups (Figure 4D). Although pre-depolarization did not differ significantly (Δ: 0.06 mV, HDI: [−0.81, 0.93]), ripple-depolarization was substantially lower in PCs than NPCs (Δ: −2.30 mV, HDI: [−3.70, −0.89]; Figure 4E). However, neither R_in_ nor θ_mean_ differed significantly between PCs and NPCs (Δ: −12.19 MΩ, HDI: [−45.52, 23.46]; Δ: −0.88 mV, HDI: [−3.03, 1.20]; Figure 4G). Further examining the combined effect of all variables using the binomial regression model fit on both variable types (Figure 2I) and decomposing their contributions (Figure 4G), we found that ripple-depolarization provided by far the largest contribution to the observed ΔPR between PCs and NPCs (ΔPR contribution: −8.81 %, HDI: [−10.03, −7.63]; Figure 4J), followed by the unexplained residuals (ΔPR contribution: −2.03 %, HDI: [−3.47, −0.58]) and θ (ΔPR contribution: −0.98 %, HDI: [−1.50, −0.47]). Both pre-depolarization (ΔPR contribution: 0.14 %, HDI: [0.02, 0.25]) and R_in_ (ΔPR contribution: −0.10 %, HDI: [−0.67, 0.43]) on the other hand contributed relatively little to the observed ΔPR. Thus, the lower ripple participation of PCs was predominantly associated with their weaker ripple-depolarization rather than systematic differences in intrinsic excitability.

### Post-ripple rebound is associated with hyperpolarization of CA1 pyramidal cells

In addition to cellular diversity, hippocampal ripple diversity has been increasingly recognized.^18–21^ We next asked how CA1 pyramidal cells respond differently to distinct ripple subtypes. To identify what signal features differentiated ripples most, we performed principal component analysis (PCA) on the z-scored ripple waveforms in the LFP. The first principal component, which explained 16.6% of the variance, showed its strongest effect in a period following the ripple peak (Figure 5A, B, black arrow). We next tested if this component was related to the previously measured post-ripple rebound and found that they were strongly correlated within each cell (β: 8.86, HDI: [8.52, 9.19]; Figure 5C). This suggests that post-ripple rebound is one of the defining features distinguishing ripples from one another, motivating us to further investigate its relationship to other ripple features. It has been shown, for example, that ripple amplitude and frequency are both positively correlated with sharp-wave amplitude and the magnitude of a current-source-density (CSD) sink in the SR.^19^ Similarly, we found that post-ripple rebound was also positively related with both ripple amplitude (β: 0.16, HDI: [0.13, 0.18]) and frequency (β: 0.17, HDI [0.14, 0.19]; Figure 5D). Therefore, the post-ripple rebound therefore adds to the LFP and CSD features that distinguish ripple subtypes.

**Figure 5:**
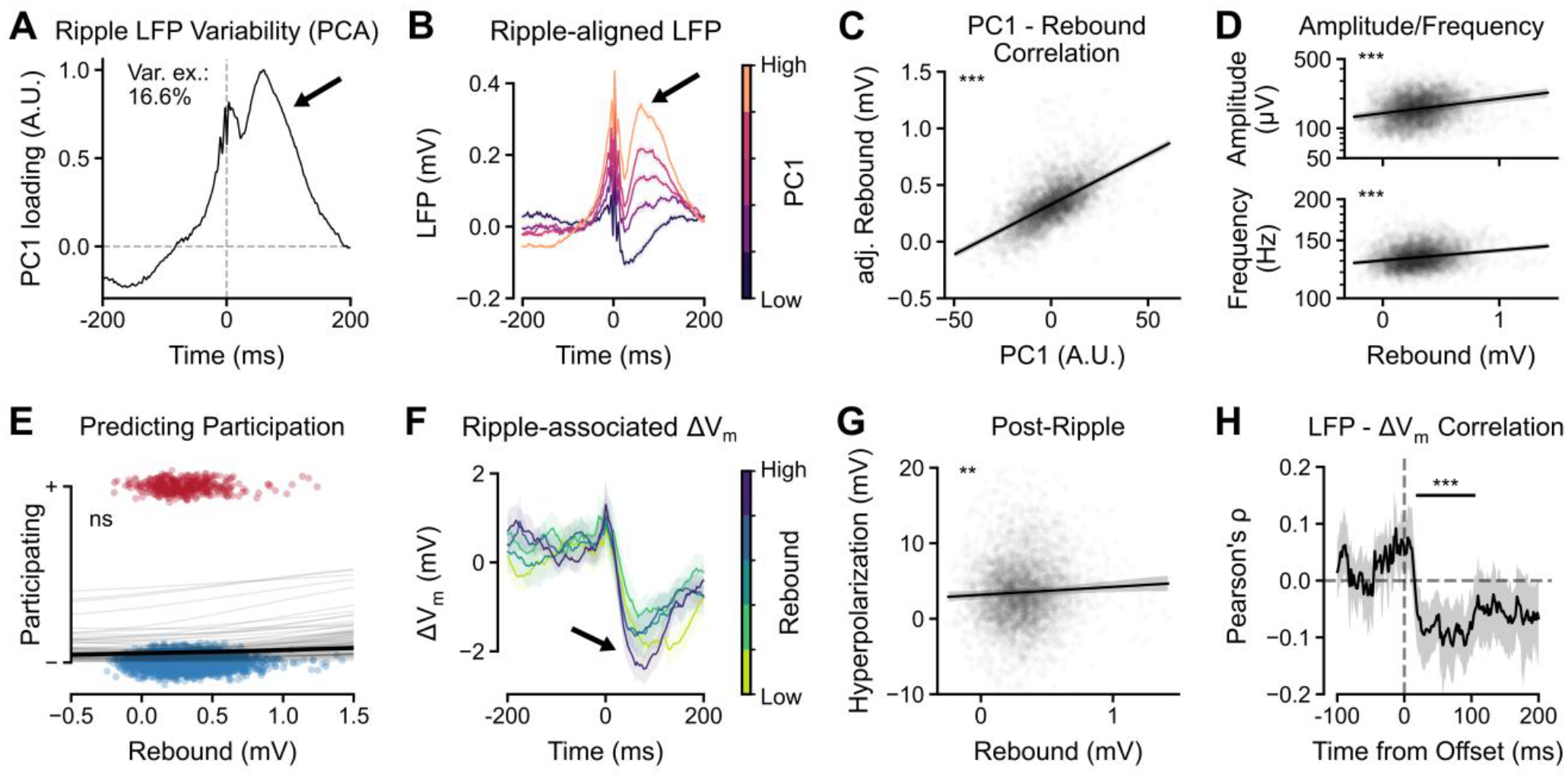
Post-ripple rebound associated with post-ripple hyperpolarization. **(A)** First principal component (PC1) loading extracted using principal component analysis (PCA) performed on the z-scored ripple waveforms aligned to ripple peak. PC1, the axis capturing the most ripple-to-ripple variability, explained 16.6% of the total variance (Var. ex.). Post-ripple rebound is highlighted with a black arrow. **(B)** Ripple-aligned LFP waveforms across PC1 quintiles, aligned to the ripple peak, showing the strong increase of post-ripple rebound (highlighted with a black arrow) with increasing PC1. **(C)** Correlation of PC1 and post-ripple rebound estimated using a linear mixed-effects model. Rebound values were adjusted for the random effect (adj. Rebound). Black line represents median posterior prediction with the 95% highest density interval highlighted in grey. **(D)** Correlation of post-ripple rebound, and ripple amplitude (top) and frequency (bottom) estimated using a linear mixed-effects model fitted on logarithmic transformed values. Amplitude and frequency values were adjusted for the random effect and plotted on a logarithmic axis. Black line represents median posterior prediction with the 95% highest density interval highlighted in grey. **(E)** Relationship between post-ripple rebound and the probability of participation. Points show individual ripple events and curves show cell-specific participation functions estimated using hierarchical logistic regression with both random intercepts and random slopes. No significant association could be found in any cell. **(F)** Ripple-associated ΔV_m_ (mean±SEM) across post-ripple rebound quintiles, showing increased post-ripple hyperpolarization (black arrow). **(G)** Correlation of post-ripple rebound, and post-ripple hyperpolarization estimated using a linear mixed-effects model. Post-ripple hyperpolarization values were adjusted for the random effect. Black line represents median posterior prediction with the 95% highest density interval highlighted in grey. **(H)** Time-resolved Pearson correlation coefficient between LFP trace and ΔV_m_, aligned to ripple offset. Shading indicates the 95% confidence interval, bootstrapped across cells. Pointwise p-values were corrected for multiple comparisons using cluster-correction. Significant clusters are marked with a horizontal bar. Statistical evidence is denoted by asterisks according to the highest credible level whose highest density interval excludes the null (*, 95%; **, 99%; ***, 99.9%; ns, otherwise).

To determine how individual CA1 pyramidal cells respond to ripples with different rebound amplitudes, we first investigated whether participation varied accordingly but were unable to identify a clear relationship between the two (β: 0.07, HDI: [−0.07, 0.21]; Figure 5E). Examining the ΔV_m_ response of cells to ripples with different rebound amplitudes, we found that larger rebounds were associated with greater post-ripple hyperpolarization (β: 0.22, HDI: [0.05, 0.38]; Figure 5F, G). To test whether the two signals are also correlated on a similar timescale, we aligned the LFP and ΔV_m_ traces to ripple offset and calculated the Pearson correlation coefficient across ripples at each timepoint. After applying a cluster-based correction method for multiple comparisons (see Methods), we identified a significant time window (20 – 105 ms after ripple offset), in which the LFP and ΔV_m_ traces were negatively correlated (Figure 5H). This indicates that ripples with a larger positive LFP rebound were associated with stronger post-ripple hyperpolarization at the single-neuron level.

## Discussion

This study investigated what cellular factors determine whether individual CA1 pyramidal cells are recruited to participate in spiking during hippocampal ripples. At the level of individual ripple events, participation was strongly associated with ripple-related membrane-potential depolarization. However, substantial differences in participation propensity remained between cells. At the cell level, differences in participation propensity were best explained by the combined influence of ripple-associated membrane-potential dynamics and intrinsic excitability, with neither factor alone sufficient to account for the observed variance. The relative balance of these two factors differed across cell classes. Superficial cells participated more frequently than deep cells, primarily because they received stronger ripple-associated depolarization, whereas differences in intrinsic excitability acted in the opposite direction and partially counterbalanced this advantage. In contrast, place cells showed lower overall ripple participation than non-place cells, a difference that was mainly attributable to weaker ripple-associated depolarization rather than to systematic differences in intrinsic excitability. Together, these findings suggest that ripple recruitment is determined by an interplay between the momentary synaptic input a neuron receives around ripples and its propensity to transform that input into spike output.

Previous studies have similarly shown that spiking of CA1 pyramidal cells during individual ripples is associated with concurrent depolarization and reflects a competition between excitatory and inhibitory synaptic drive. Our results confirm this relationship at the level of individual SWR events: fluctuations in pre-ripple and ripple-associated depolarization were informative about whether a neuron would participate in a particular ripple. However, cell-to-cell heterogeneity (H_cell_) also contributed substantially to predictive performance. To account for between-cell differences, we next investigated cell-intrinsic properties, focusing on input resistance and AP threshold. Input resistance is a natural candidate, since it determines the membrane-potential response produced by a given synaptic current. When input resistance and AP threshold were used to predict participation ratio without membrane-potential variables, higher input resistance was associated with greater participation propensity. However, this association was strongly reduced once ripple-associated depolarization was included in the model, indicating that the effect of input resistance on participation runs largely through its influence on membrane-potential responses.

We used the model-estimated AP threshold throughout this study, rather than AP threshold measured from current-injections. This choice was motivated in part by a practical constraint: fluctuations in access resistance, a common challenge during intracellular patch-clamp recordings in head-fixed, running mice, could not always be perfectly compensated for in real time by bridge balance, introducing variability into current-injection-based threshold measurements that was unrelated to genuine differences in cell excitability. More fundamentally, the model-estimated θ offers a conceptual advantage: it is derived from the same spontaneous, ripple-associated ΔV_m_ traces analysed throughout this study, rather than from a separate current-injection protocol subject to its own distinct technical confounds. To estimate the AP threshold, we fitted a phenomenological spike-generation model to spontaneous membrane-potential dynamics in which threshold was represented by a distribution. This yielded two interpretable parameters: θ_mean_, describing the location of the inferred threshold distribution, and θ_SD_, describing its apparent variability. As expected, a more depolarized θ_mean_ was associated with lower participation propensity (Figure 2G). In contrast, greater θ_SD_ was associated with higher participation propensity. This result suggests that increased threshold variability raises the likelihood that threshold falls sufficiently low for some ripple events to permit firing. Apparent threshold variability can arise from several sources. AP threshold is itself dynamic and can depend on recent membrane-potential history, sodium-channel availability, preceding spiking and the rate of depolarization.^26,27^ In addition, fluctuations in unobserved synaptic conductance or dendritic inputs may appear in a phenomenological model as variability in AP threshold.

CA1 pyramidal cells form functionally distinct sublayers: DCs and SCs differ in firing and bursting properties, spatial coding and their coupling to hippocampal oscillations.^16,23^ Previous work demonstrated that SCs receive stronger net depolarizing drive during ripples, whereas DCs can experience stronger inhibition.^16^ Our results are consistent with this distinction but additionally show that differences in intrinsic excitability can oppose differences in synaptic drive. SCs showed stronger ripple-associated depolarization, which promoted their recruitment, whereas differences in inferred AP threshold acted in the opposite direction. Their greater participation therefore did not arise simply because superficial cells were intrinsically more excitable. Instead, participation reflected the net effect of multiple cellular factors that could either reinforce or counteract one another. This framework may help reconcile observations that DCs exhibit greater firing or bursting propensity under some conditions while SCs can nevertheless be preferentially recruited during ripples.^16,23^

A different combination of factors distinguished PCs from NPCs. PCs participated, on average, in fewer ripples, and this difference was associated primarily with weaker ripple-associated depolarization rather than with systematic differences in the intrinsic excitability measures examined here. A substantial portion of this difference, however, was left unexplained. This residual difference could reflect cellular properties not captured by our measurements, differences in excitatory or inhibitory connectivity, or, importantly, the selective recruitment of place cells according to the content represented by individual ripples.

At first sight, reduced global participation of PCs may appear surprising because replay is classically defined by the reactivation of place-cell sequences. However, participation probability and replay selectivity describe different properties. Replay does not require every PC to participate frequently across ripples; rather, specific subsets of place cells must be recruited during particular events according to the represented trajectory or experience. Lower average participation of PCs could therefore coexist with strong recruitment during the subset of ripples representing the relevant location, trajectory or experience. Such selective recruitment could preserve the specificity of replayed representations and limit interference between competing spatial ensembles. This interpretation remains speculative because we could not decode the representational content of individual ripple events. An important future test will therefore be to determine whether low-participation PCs are depolarized only when they are part of a specific decoded neuronal replay sequence and hyperpolarized otherwise.

Our final results highlight an additional dimension of hippocampal ripple heterogeneity. The post-ripple LFP rebound emerged as the most prominent feature differentiating individual ripple events, with rebound amplitude covarying strongly with ripple amplitude and frequency. This finding echoes earlier attempts to classify ripple subtypes using different approaches.^19,20^ Ramirez-Villegas et al.^20^ clustered ripple waveforms recorded in urethane-anesthetized macaques and identified “classical” ripples to have a higher ripple frequency compared to “non-classical” ripples. Castelli et al.^19^ classified ripples based on current-source density (CSD) profiles and found Rad^sink^ ripples (CA3 input-driven) to have a higher ripple frequency compared to LM^sink^ ripples (EC input-driven). Ripple frequency therefore separates subtypes across three independent approaches: waveform clustering, laminar current profile, and post-ripple rebound, suggesting that these approaches converge on a shared underlying distinction. Importantly, larger LFP rebounds were associated with stronger post-ripple hyperpolarization of CA1 pyramidal cells. Previous studies have shown that ripples are followed by membrane-potential hyperpolarization and suppression of pyramidal-cell firing^13^, and that the magnitude of post-ripple hyperpolarization covaries with sharp-wave amplitude.^14^ Our results extend these observations by directly linking a concurrent extracellular feature of the ripple waveform to the post-ripple membrane-potential response of individual CA1 pyramidal cells.

What are the mechanisms that could be responsible for post-ripple hyperpolarization? Inhibition is central to ripple generation, with parvalbumin-expressing interneurons providing rhythmic perisomatic inhibition during ripples, but the mechanisms that determine ripple termination and subsequent suppression remain incompletely understood. Stronger population excitation could recruit correspondingly stronger feedback inhibition, eventually overcoming recurrent excitation and contributing to ripple termination. CCK+ interneuron subtypes are known to preferentially fire outside of ripples during quiescence,^16,28^, while slowly acting dendrite-targeting interneurons of the neurogliaform or Ivy-cell family may contribute to inhibitory control around and following ripple termination.^29^ The association between rebound magnitude and cellular hyperpolarization is compatible with such mechanisms. Future experiments involving direct recordings or selective manipulation of identified inhibitory populations will be required to determine which circuits generate the post-ripple hyperpolarization and whether they actively contribute to terminating or separating ripple events.

## Acknowledgments

This project acknowledges funding from the Swiss National Science Foundation (PR00P3_208598, PR00P3_208598/2, TMPFP3_210606, X.Z.), the Synapsis Foundation (Grant Agreement 2022-CDA02, X.Z.), and The URPP AdaBD. We thank David Dupret, Manfredi Castelli, and Peter Rupprecht for their insightful discussions and valuable feedback on earlier versions of this manuscript. Special thanks to Vladimirs Dobrovolskis for assistance with brain slicing and confocal imaging, and Dubravka Dujmovic Göckeritz for technical support. We also appreciate Stefan Giger, Martin Wieckhorst, Hansjörg Kasper, and Frank David for their contributions to constructing the setup and providing IT support.

## Author contributions

X.Z. conceived the project and performed experiments. X.Z., R.A. and T.L. analysed data. X.Z. and R.A. wrote the paper. P.J. and F.H. provided conceptual input and resources. All authors jointly revised the paper.

## Declaration of interests

P.J. has an industrial collaboration with Leica Microsystems and is member of the advisory board of Neuron. The other authors declare no competing interests.

## Declaration of the use of generative AI

During the preparation of this work, the authors used ChatGPT (OpenAI) to assist with language editing and debugging of analysis code. After using this tool, the authors reviewed and edited the content as needed and take full responsibility for the content of the published article.

## Methods

### Key Resources Table

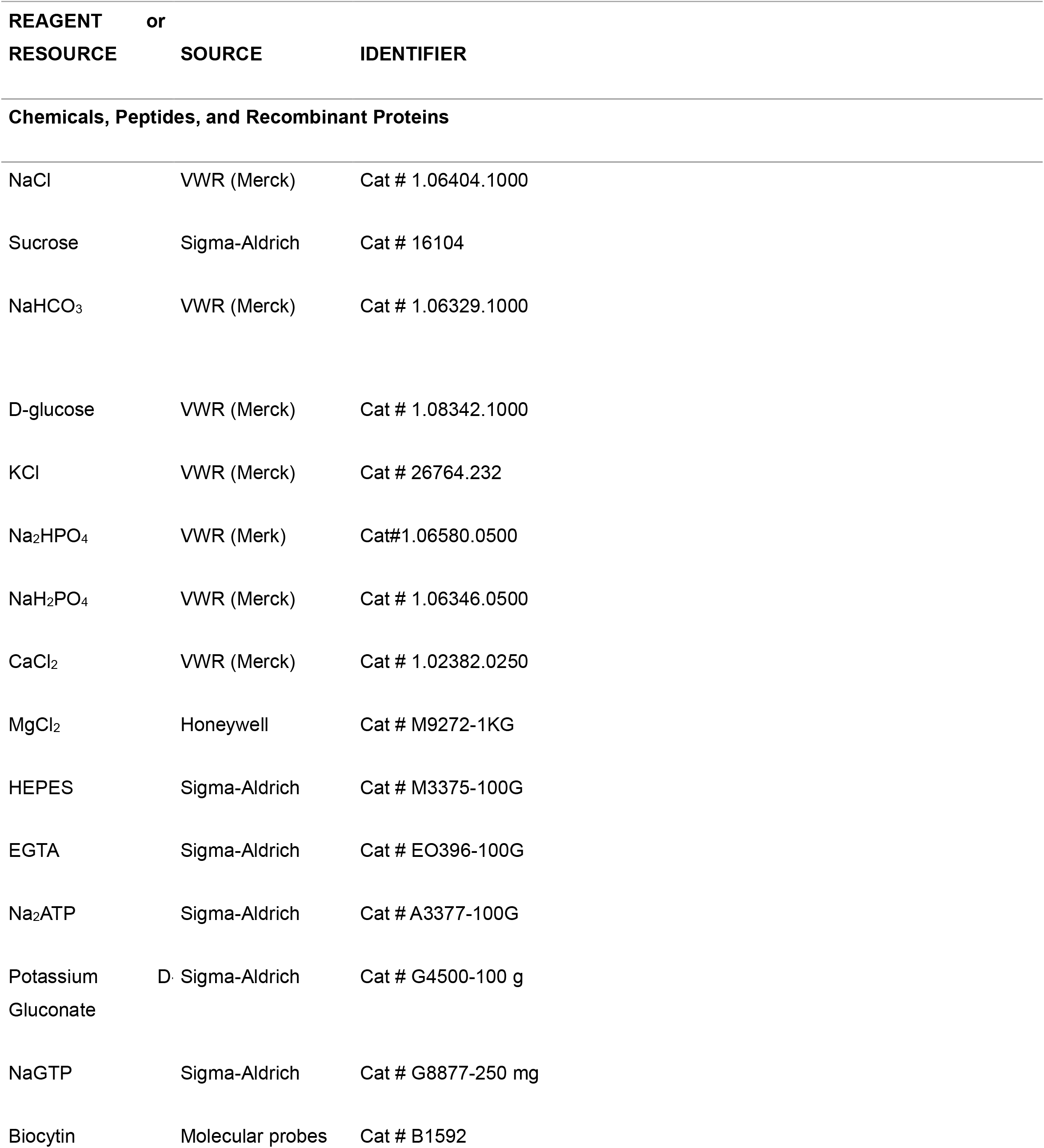
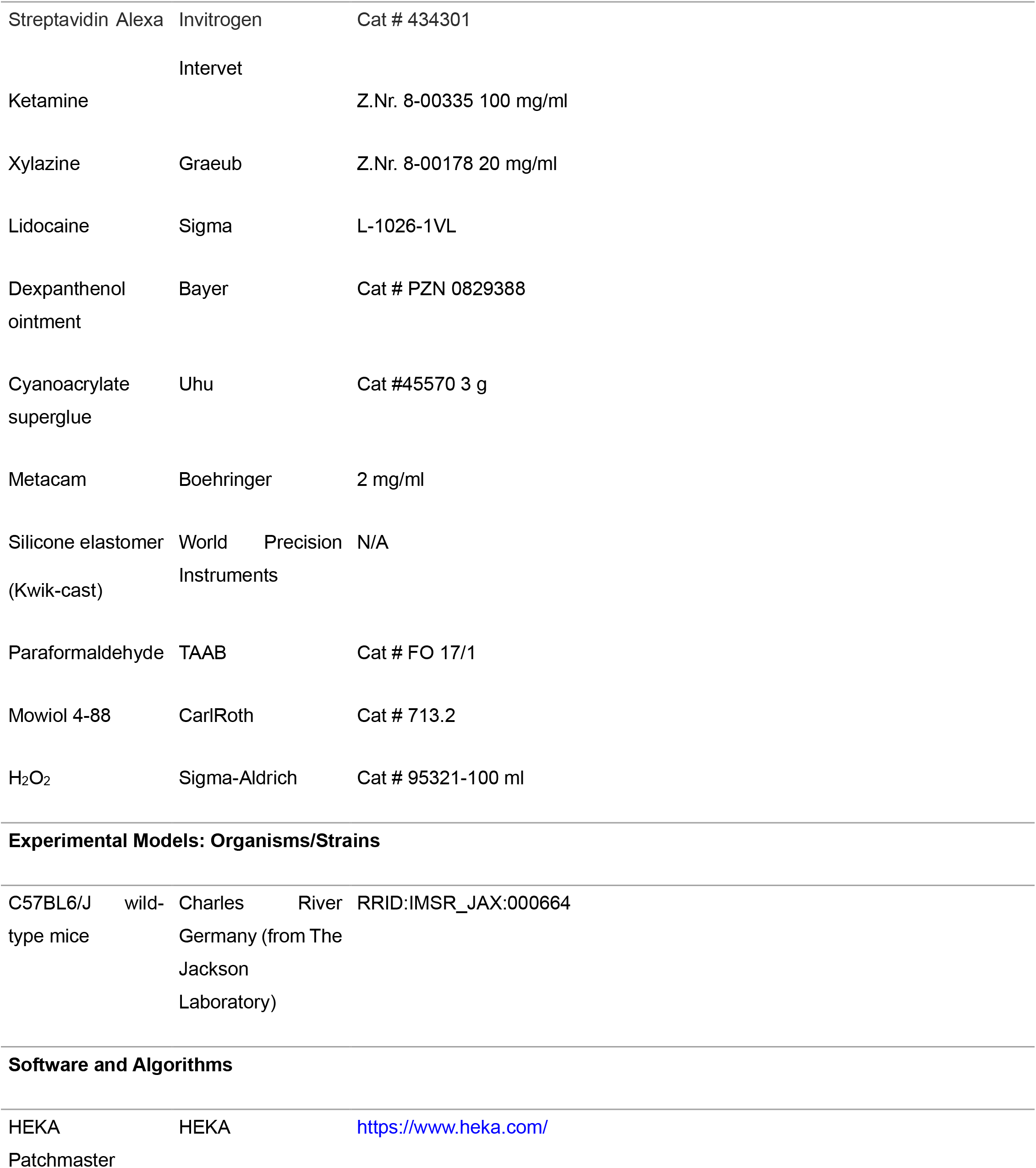
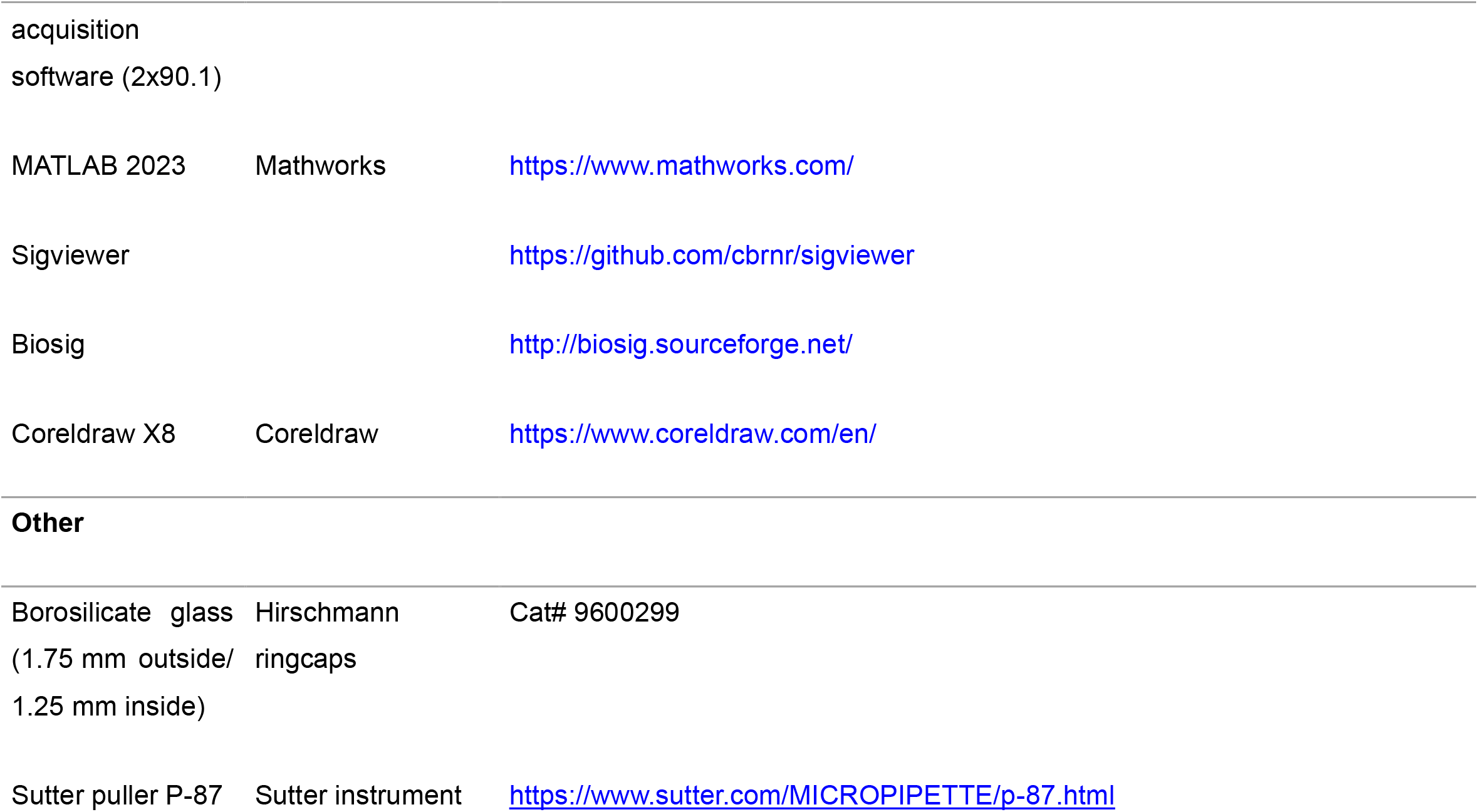

### Resource Availability

#### Lead Contact

Further information and requests for resources should be directed to and will be fulfilled by the Lead Contact, Xiaomin Zhang.

#### Materials Availability

This study did not generate new unique reagents.

#### Data and Code Availability

Original data, analysis programs, and computer code were stored in the scientific repositories of the Brain Research Institute of the University of Zürich and are available upon reasonable request.

### Experimental Model and Subject Details

#### Animal experiments

Whole-cell patch-clamp recordings *in vivo* were performed in 35-to 63-day-old C57BL/6 mice (RRID:IMSR_JAX:000664). Animals were housed under a reversed light cycle (dark: 7:00 am – 7:00 pm, light: 7:00 pm – 7:00 am). For experiments, both male and female animals were used. All experiments were carried out in strict accordance with institutional, national, and Swiss guidelines for animal experimentation, and approved by the Zürich cantonal authority (ZH 019/2022).

#### Surgery and animal training

Head-bar implantation and craniotomy were performed under isoflurane anaesthesia (5% for induction, 1.5-2% for maintenance), supplemented with local lidocaine anaesthesia (Astra Zeneca), as described before.^30,32^ Oxygen was provided throughout the procedures. EMLA cream was applied to the ear bars of the stereotaxic frame to minimize pressure sensation on the external auditory meatus. Dexpanthenol ointment (Bayer) was applied to the eyes to prevent corneal drying. The skull was gently abraded using a dental drill and cleaned with saline solution (0.9% NaCl). A custom-made steel head-bar was affixed to the skull using Scotchbond (3M) and stabilized with Charisma dental cement (Kulzer). Mice were kept on a heating pad until fully awake. Analgesia was ensured with intraperitoneal administration of 50 mg kg-1 buprenorphine (Sanofi-Aventis) twice daily for three days post-surgery. Mice received a nutritionally enriched diet following head-bar surgery. After a one-week recovery period, running wheels were introduced into the home cage. Mice underwent citric acid water restriction (2%, Sigma) and daily handling by the experimenter for approximately 10 minutes over 3-4 days. Once accustomed to water restriction and handling, mice were trained to run on a linear treadmill for a water reward lasting 45 minutes per day. Water rewards were dispensed when the animal covered a distance of 180 cm. Training sessions continued for 7-10 days until consistent daily running distances were achieved over three consecutive days. The day before recording, two small craniotomies (~0.5 mm diameter each) were gently drilled at coordinates approximately −1.8 mm and −2.3 mm anterior-posterior (for whole-cell recording and local field potential, respectively, measured from bregma) and ~1.2 mm medio-lateral (measured from midline). The dura mater remained intact, and craniotomies were sealed with silicone elastomer (Kwik-cast, World Precision Instruments).

#### Behavioural set-up

Custom-built linear treadmills were used for both behavioural training and electrophysiological recordings. The treadmill belts were 180/135 cm in length and constructed from velvet fabric (McMaster-Carr). Each belt was divided into three equal segments, each marked by distinct tactile cues: glue spines, Velcro tabs, and zigzag glue tracks, as previously described.^30,32^ Water rewards were delivered via a syringe pump, triggered by a TTL pulse upon entry into the glue spine segment. Licking behaviour was monitored using a custom-designed piezoelectric sensor lick port that detected pressure deflections corresponding to tongue contact. Animal position along the belt was tracked using an infrared (IR) reflective sensor (RS Components) in combination with reflective tape placed at the end of each textured segment, which also served to trigger reward delivery. Locomotor speed and position were continuously measured using an incremental rotary encoder mounted on one of the treadmill wheel axes. An Arduino-based interface was employed to generate analog signals representing velocity, position, texture segment, and licking events. These signals were routed to the analog input channels of a Heka amplifier to enable precise synchronization with electrophysiological recordings

#### *In vivo* electrophysiology

In vivo whole-cell patch-clamp recordings were obtained from CA1 pyramidal cells (PCs) in the dorsal hippocampus, following previously established protocols.^30,32^ Patch and LFP pipettes were pulled from borosilicate glass capillaries (outer diameter: 1.75 mm; inner diameter: 1.25 mm) using a horizontal puller (P-87, Sutter Instrument). Local field potential (LFP) electrodes (3–5 MΩ) were filled with artificial cerebrospinal fluid (ACSF) and mounted on a second, custom-modified micromanipulator, angled 20° posteriorly relative to bregma. The LFP electrodes were targeted to the CA1 pyramidal cell layer, confirmed by the presence of sharp wave-ripples. Whole-cell patch pipettes (9–12 MΩ) with a long taper and were filled with an intracellular solution containing (in mM): 130 K-gluconate, 2 KCl, 10 HEPES, 2 MgCl_2_, 2 Na_2_ATP, 0.3 NaGTP, 18 sucrose, 10 EGTA, and 0.3% biocytin for post hoc morphological analysis. The pH was adjusted to 7.3 with KOH. To minimize clogging during cortical penetration, whole-cell electrodes were advanced with positive pressure of 500–600 mbar. Upon reaching a depth of ~800 µm, pressure was reduced to ~20 mbar. Putative somatic contact was identified by a consistent increase in pipette resistance. Gentle suction was then applied to establish whole-cell access. Access resistance was monitored using voltage test pulses (+50 mV and −10 mV); recordings were discarded if series resistance exceeded 100 MΩ. Following bridge balance compensation, a series of step currents (−100 to +400 pA) was injected to assess input resistance and maximal action potential firing rate. All recordings were conducted in whole-cell current-clamp mode with zero current injection using a HEKA EPC double amplifier. Pipette capacitance was minimized, and access resistance was compensated via the amplifier’s bridge-balance circuit. Signals were low-pass filtered at 10 kHz and digitized at either 10 or 25 kHz using Patchmaster software (v2×90.1, HEKA). At the end of each recording, the patch pipette was slowly withdrawn to form an outside-out patch, allowing for assessment of seal quality and reliable biocytin labelling of CA1 PCs.

#### Neuron labelling and confocal imaging

After the recording, animals were transcardially perfused with 20 ml of 0.1 M phosphate buffer (PB), followed by 20 ml of 4% paraformaldehyde (PFA). Brains were carefully extracted and post-fixed by immersion in 4% PFA overnight at 4°C. Subsequently, brains were transferred to phosphate-buffered saline (PBS) for long-term storage. 150-μm-thick coronal brain slices were prepared and washed three times in 0.1 M PB on a shaker at room temperature. Slices were then incubated overnight at 4°C in antibody solution containing Streptavidin conjugated to Alexa Fluor 488 (1:500, Invitrogen) diluted in 0.1 M PBS with 1% Triton X-100. The next day, slices were washed three times for 10 minutes each in 0.1 M PBS on a shaker at room temperature in the dark. Finally, slices were mounted onto microscope slides, dried briefly, coverslipped with Mowiol mounting medium (Roth), and stored at 4°C until confocal imaging.

The image acquisition was performed on the Olympus Fluoview FV1000 confocal microscope, equipped with a UPLSAPO 10x and 20x, NA0.40 objective. The acquisition process was controlled through Olympus Fluoview v4.2x software, with pixel dwell time set between 8.0 µs/pixel in fast mode, with an image capture speed of approximately 5.1 seconds. The resolution was configured at 800×800 pixels, providing high enough detail. Fluorescence imaging was carried out using GFP channels, which was excited at 488 nm. The laser power was optimized to prevent photobleaching while ensuring sufficient signal intensity. The gain and offset settings were also customized for each channel to optimize the signal-to-noise ratio. The images were saved in TIF and OIF formats to preserve quality for more detailed inspection and post-processing on ImageJ software.

Confocal images with a singular Biocytin-labelled soma and a distinguishable pyramidal layer were analysed using MATLAB (2023b; MathWorks). Using a custom graphical user interface (GUI), the centroid of the soma was manually selected, and the borders of the pyramidal layer were manually drawn and interpolated using splines. The distance of the soma centroid from the stratum pyramidale/ radiatum and stratum pyramidale/ lacunosum molecular borders were then computed as the minimal Euclidean distance. The radial depth was inferred as the percentage of the distance from the stratum pyramidale/ radiatum over the total distance.

### Data analysis

#### Spike detection, identification, and removal

The 0.6 ms centre differential of the raw V_m_ recording was taken, similar to the previous work.^14^ Positive- and negative-going windows were taken as epochs where the centre differential was respectively greater than 5 V/s and lower than −2.5 V/s. Spike windows were then detected as positive-going windows preceding the negative-going window within 1.5 ms. Spike peaks were detected as the maximum V_m_ within the detected window. The windows were then expanded by [−8 1.2] ms relative to the detected window to get the spike window. If consecutive spike windows were overlapping, the minimum V_m_ between the two windows’ peaks were identified, and their respective off-and on-set were then respectively redefined as the 0.04/0.10 ms preceding and following the minimum. By doing so, the subthreshold Vm fluctuations during spike bursts could maximally be preserved, such as during plateau potentials.

Spike slope, threshold, and amplitude were defined for each spike window detected. Maximal spike slope was measured as the maximum 0.6 ms centre differential within the spike window. AP-threshold was measured as previously described.^31^ The threshold slope was taken as 10 V/s if the maximal slope is greater than 30 V/s or a third of the maximal slope if it is smaller than 30 V/s. Spike onset was defined as the timestamp when the centre differential crosses the threshold slope. Spike amplitude was measured as the difference between the peak amplitude and the spike threshold. Spikes below 30 mV were excluded from the analysis, and the remaining were classified as action potentials (AP).

The V_m_ traces were linearly interpolated across all spike windows to reveal the subthreshold V_m_ (SubV_m_) fluctuations. The SubV_m_ was zero-phase low-pass filtered using a 4th order Butterworth filter with a 400 Hz cutoff, then downsampled to 2500 HZ. Sharp edges from linear interpolation could therefore be minimized to preserve the smooth SubV_m_ fluctuations. The LFP traces were zero-phase, high-pass filtered using a 4th order Butterworth filter with a 0.3 Hz cutoff to remove the slow drift components from the recordings. A zero-phase, low-pass, 8th order Butterworth filter with a 400 Hz cutoff, then a downsample to 2500 Hz were applied. Movement epochs were differentiated from immobility epochs by thresholding the velocity data of the linear treadmill. A 0.5 cm/s threshold was used, and movement epochs were merged if they were found within 3 s.

#### Quantification of intrinsic parameters during current injections

Passive and active membrane properties of recorded cells were obtained by injecting 1 s current pulse starting from −100 pA to 400 pA with 50 pA increments. Resting membrane potential was obtained by taking the median Vm across the 100 ms windows prior to the current injections. Rheobase was defined as the minimum current injection required to elicit at least one AP. The relative AP threshold was defined as the V_m_ at spike onset of the first AP at Rheobase subtracted by the resting membrane potential. Input resistance (R_in_) was measured as the slope of the robust linear fit between the injected currents and the elicited median V_m_ during current injections that did not trigger APs.

#### Sharp-wave ripple detection and characterization

Sharp-wave ripples were detected from the CA1 LFP during immobility. Movement epochs were identified from treadmill velocity using a threshold of 0.5 cm/s, and ripple detection was restricted to the complementary stationary periods. Stationary segments shorter than 1 s were excluded. The LFP was band-pass filtered between 100 and 250 Hz and transformed using the Hilbert transform. The magnitude of the resulting analytic signal was used as the ripple-band envelope. To reduce sensitivity to large-amplitude outliers, detection thresholds were defined using robust estimates of the centre and scale of the envelope. The centre was given by the median envelope magnitude and the robust scale by

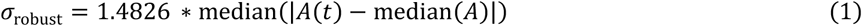

where *A*(*t*) denotes the ripple-band envelope. Candidate ripple intervals were first identified as contiguous periods during which the envelope exceeded 3 times *σ*_robust_. A candidate was retained only if its envelope additionally exceeded 8 times *σ*_robust_ at least once within the interval. Candidate intervals separated by less than 20 ms were merged. Events shorter than 20 ms or longer than 500 ms were excluded.

For each candidate event, ripple phase was obtained from the analytic signal and unwrapped across the event. The number of oscillatory cycles was calculated from the total phase progression divided by 2π, and events containing fewer than 1.8 cycles were excluded. Ripple frequency was calculated as the number of cycles divided by ripple duration. The ripple peak was defined as the time of the maximum ripple-band Hilbert envelope within the detected interval. Final ripple detections were manually validated using a custom graphical interface displaying the raw LFP together with a time-frequency representation spanning 80–300 Hz. Candidate events were accepted or rejected following visual inspection, and only accepted events were included in subsequent analyses. Ripple amplitude was measured independently as the maximum analytic amplitude within each accepted ripple after 100–250-Hz fourth-order Butterworth band-pass filtering of the LFP. Inter-event interval of a specific ripple was defined as the minimum time to either of its neighbours. Rebound amplitude was defined as the mean LFP of the 200 ms period following ripple offset.

#### Ripple-associated membrane-potential dynamics

For each accepted ripple, an event-specific membrane-potential baseline (Baseline V_m_) was estimated from ripple-free SubVm samples between 5s before and after ripple onset and offset respectively. To prevent contamination by nearby ripple events, ripple intervals were excluded together with 200-ms padding on either side. Baseline V_m_ was defined as the median SubVm within the remaining baseline samples. Subtracting this value from the SubVm trace yielded the ripple-triggered SubVm change (ΔV_m_). Pre-ripple depolarization was subsequently defined as the mean ΔV_m_ between 100 and 25 ms before ripple onset. To isolate ripple depolarization the 95^th^ percentile of the ΔV_m_ samples within the detected ripple interval was subtracted by the pre-ripple depolarization. Post-ripple hyperpolarization was quantified as the negative of the 5^th^ percentile of the ΔV_m_ within 200 ms following ripple offset, such that larger positive values indicate stronger hyperpolarization.

#### Event-level ripple participation

An individual ripple was classified as participating (P+) for a given neuron when at least one AP onset occurred between the detected ripple onset and offset. Events without an AP during this interval were classified as non-participating (P-). To determine whether event-to-event fluctuations in membrane potential predicted participation independently of persistent differences between neurons, Pre and Ripple depolarization were mean-centred within each cell:

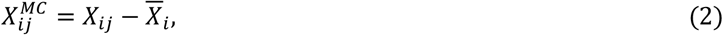

where (i) indexes cells and (j) indexes ripple events. Ripple participation was subsequently modelled using a hierarchical logistic regression,

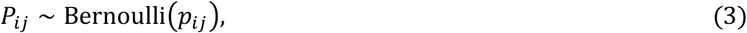

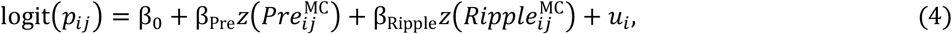

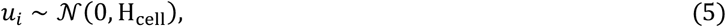

with Pre and Ripple denoting pre- and ripple depolarization respectively and u_i_ a cell-specific random intercept. Continuous predictors were standardized before model fitting. Mean-centring isolates within-cell variation, such that the fixed effects quantify whether a given neuron was more or less likely to participate when its membrane-potential response differed from its own average response and the random effect isolates the cell-to-cell differences in participation propensity (H_cell_).

The discriminative performance of this model was quantified using a receiver-operating-characteristic curve (ROC) and the area under the curve metric (AUC). AUC was calculated separately for posterior draws to propagate uncertainty in the fitted model. The contributions of Pre depolarization, Ripple depolarization, and cell identity to the improvement in AUC were additionally decomposed using Shapley values (see Shapley decomposition).

#### Shapley decomposition

To quantify the contribution of individual predictors to model-derived outcomes, we used Shapley decomposition, where the contribution of each predictor is quantified by a Shapley value. These have the property that their sum together with the null prediction (meaning using the overall mean for each predictor) give the prediction using all the data. For each posterior draw, predictions were generated for every possible coalition of predictors. Predictors included in a coalition were retained at their observed values, whereas excluded predictors were set to their reference value, corresponding to the mean of the respective predictor. For a given quantity of interest, a value *v*(*S*) was calculated for each predictor coalition S. The Shapley value *ϕ* of predictor j was then defined as its average marginal contribution across all possible coalitions,

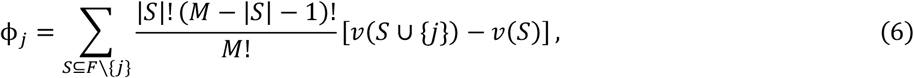

where F is the full set of predictors and M is the number of predictor components. This procedure accounts for dependence and shared explanatory information between predictors by averaging the contribution of each predictor over all possible orders in which predictors can enter the model. All decompositions were performed on the probability scale after applying the inverse-logit transformation. Depending on the analysis, *v*(*S*) represented either the AUC, the between-cell variance in predicted participation probability or the difference in mean predicted participation probability between two cell groups. The decomposition was calculated independently for every posterior draw, and Shapley contributions are therefore reported as posterior distributions with corresponding 95% highest-density intervals.

#### Participation Ratio

For each cell, participation ratio (PR) was defined as

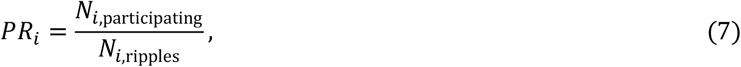

where *N*_*i*,participating_ is the number of accepted ripples containing at least one AP and *N*_*i*,ripples_ is the total number of accepted ripples recorded from cell (i). To estimate the participation propensity of individual neurons while accounting for the number of ripple events recorded from each cell, participating and total ripple counts were modelled using a hierarchical binomial model,

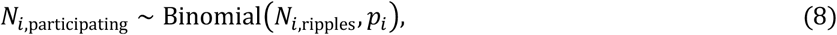

with a cell-specific random intercept on the logit scale. The posterior median of the resulting cell-specific participation probability was used to rank neurons. Based on this rank, cells were divided into five equally sized participation-rate groups for visualization. For cell-level analyses, mean Pre and Ripple depolarization were calculated by averaging the corresponding event-level quantities across all accepted ripples recorded from each neuron.

#### Variance explained

To quantify how much of the between-cell variation in ripple participation was captured by a cell-level model, variance explained was calculated on the probability scale. For each posterior draw, the model was used to predict the participation probability PR_predicted_ of every cell using its observed predictor values. The variance of these predicted probabilities across cells was then compared with the variance of the empirically observed participation rates (PR_observed_),

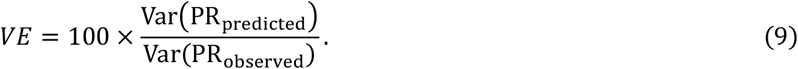

This calculation was performed independently for each posterior draw, yielding a posterior distribution of the percentage of between-cell participation-rate variance explained by the model. Variance explained is reported as the posterior median and 95% highest-density interval. Because uncertainty in the model parameters is propagated through the calculation, individual posterior draws can yield values greater than 100%.

#### Cell-level predictors of participation

Cell-to-cell variation in participation was analysed using binomial regression of participating and total ripple counts:

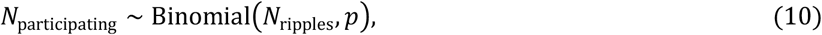

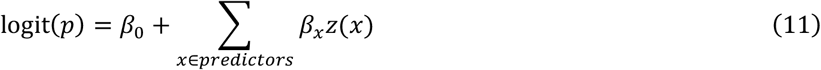

All continuous predictors were standardized before fitting. Partial-effect curves were generated by varying the predictor of interest while fixing other model predictors to their respective training-data means. Where adjusted data points were shown, partial-residual-style values were calculated on the probability scale by subtracting the full-model prediction from the observed PR and adding back the prediction attributable to the predictor of interest. The contribution of individual cellular properties to between-cell variation in participation was quantified using a Shapley variance decomposition described above.

#### Probabilistic action-potential threshold model

To estimate it from spontaneous activity, the AP threshold was modelled as a normally distributed variable characterized by a cell-specific mean threshold, θ_mean_, and threshold variability, θ_SD_:

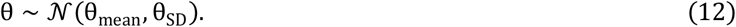

For a given membrane potential Δ*V*_*m*_, the probability that Δ*V*_*m*_ is larger than the threshold, and therefore that an AP can be generated, is given by the cumulative distribution function of this Gaussian distribution:

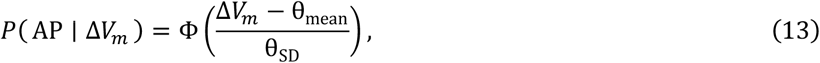

where Φ denotes the standard normal cumulative distribution function. We converted this probability into the instantaneous AP-rate by scaling it by a maximum firing rate, *λ*_max_:

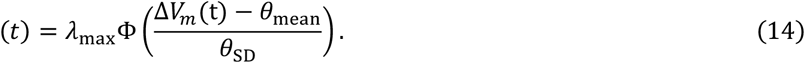

To account for the relative refractory period after each AP, *θ*_mean_ was allowed to vary over time. Each preceding AP increased the threshold an effect that decreased according to a double exponential function parametrized by a fast (*τ*_fast_ = 35ms) and a slow component (*τ*_slow_ = 500ms) such that

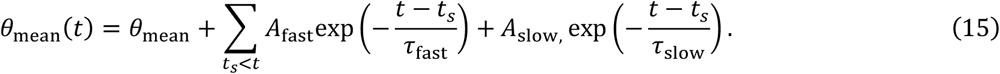

The time constants were predefined and taken from a previous study on cortical pyramidal cells.^33^ This allowed us to precompute the history effects for every timepoint, drastically reducing the computational complexity the above formula then becomes the following.

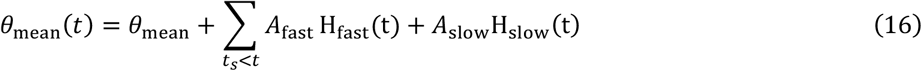

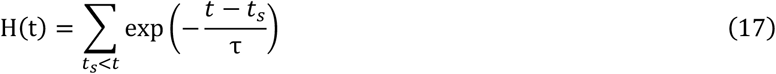

As we measured the Ripple-associated depolarization relative to a ripple-specific baseline, we aimed to emulate the baseline correction procedure on the whole SubV_m_ trace by subtracting a continuous baseline, estimated by a 10s rolling median filter, yielding a continuous Δ*V*_*m*_ trace. This means that the threshold parameters were estimated relative to the local membrane-potential baseline rather than in absolute voltage units. A 5 ms window was removed after every AP to remove spike-related afterpotentials. Furthermore, large, visually apparent artefacts were removed by visual inspection. H_fast_ and H_slow_ were calculated on the complete recording before restriction to clean intervals, using the two exponential components described above. This preserved the contribution of spikes preceding the beginning of an analysed interval. The membrane-potential trace and spike-history terms were subsequently restricted to the same clean intervals.

Rather than fitting the model to every individual time sample, the recording was summarized according to the combination of membrane potential and recent spike history. For each cell, relative membrane potential was divided into 19 bins spanning its 10^th^ to 99.9^th^ percentile. The fast and slow spike-history effects were each divided into nine bins spanning zero to their respective 99^th^ percentiles. Samples falling outside these ranges were assigned to the nearest edge bin. For each combination of membrane potential and spike history, we calculated the total amount of time spent in that bin, (*T*_*k*_), and the number of APs occurring in that bin, (*N*_*k*_). APs were assigned according to the membrane potential and spike-history values at AP onset. The resulting occupancy and spike-count distributions therefore describe the empirical AP rate as a function of membrane potential and recent spiking activity while excluding the AP waveform itself. The expected rate of APs in each bin k was obtained from the Gaussian-threshold escape rate described above,

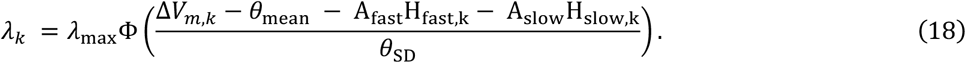

Multiplying by the occupancy T_k_ yields the expected number of spikes which we modelled as being distributed according to a Poisson distribution:

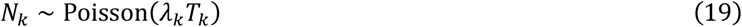

The model was hierarchical across cells, with cell-specific estimates of *θ*_mean_, *θ*_SD_, and the amplitudes of the fast and slow spike-history components A_fast_ and A_slow_. As only few neurons reached their maximum AP-rate, *λ*_max_ was estimated as a population-level parameter. Positive-valued parameters, including *θ*_SD_ and the spike-history amplitudes, were constrained to be positive by applying a softplus transform.

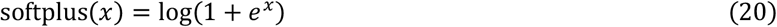

Posterior distributions were sampled using the No-U-Turn Sampler (NUTS) with four chains, 4,000 tuning iterations and 4,000 posterior draws per chain, and a target acceptance probability of 0.95. Cell-specific posterior estimates of *θ*_mean_ and *θ*_SD_ were subsequently used as measures of AP threshold and threshold variability in the cell-level participation analyses.

For visualization of membrane-potential dynamics independently of the voltage shifts associated with recent APs, ripple-aligned ΔV_m_ traces were corrected using the spike-history terms estimated by the threshold model. For each cell, the predicted history-dependent shift in mean threshold was calculated from all preceding APs as the sum of a fast and a slow exponentially decaying component, with time constants of 35 ms and 500 ms, respectively. This predicted history contribution was subtracted from the membrane-potential trace,

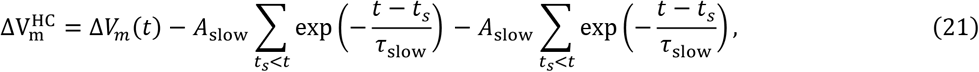

where (*t*_*s*_) denotes the time of each preceding AP.

#### Anatomical classification of CA1 pyramidal cells

For morphologically recovered neurons, radial soma position within SR was quantified from confocal images as described above. Cells with a relative distance to the SR below 0.4 were classified as superficial CA1 pyramidal cells, whereas the remaining cells were classified as deep cells. Differences in participation between superficial and deep cells were assessed using a cell-level binomial model,

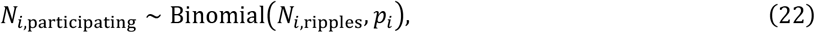

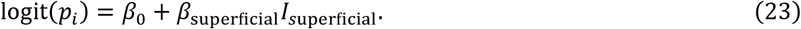

The anatomy coefficient was exponentiated to obtain the odds ratio. For descriptive interpretation, the empirical difference in mean participation rate between superficial and deep cells was additionally calculated in percentage points. Group differences in Pre and Ripple depolarization, R_in_ and θ_mean_ were evaluated using Student-t regression models with anatomical class as predictor.

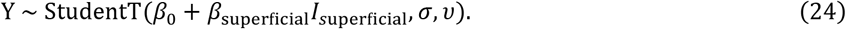

To determine how cellular properties contributed to the superficial–deep difference in participation, participation ratio was predicted using the full model fit previously and the difference between superficial and deep cells calculated. Shapley values were calculated for the contributions of Pre depolarization, Ripple depolarization, θ (both θ_mean_ and θ_SD_ together), and R_in_ to the model-predicted difference in mean participation probability between superficial and deep cells. The predicted difference of the full model to the observed empirical difference in participation ratio was taken as the part unexplained by all the factors combined. Differences in overall firing rate and bursting between superficial and deep cells were assessed from APs recorded across the full recording period. Firing rate was analysed by modelling the total number of APs per cell using a negative-binomial regression with anatomical class as predictor and the logarithm of recording duration as an offset, thereby accounting for differences in recording length. The anatomical coefficient was exponentiated and reported as a rate ratio. To quantify bursting, APs were classified as burst-associated when either the preceding or following inter-spike interval was <5ms. The number of burst-associated APs relative to the total number of APs was then modelled using a binomial regression with anatomical class as predictor, and the group effect was reported as an odds ratio.

#### Place-cell analysis

Place-cell analyses were restricted to periods in which running speed exceeded 0.5 cm/s. Individual traversals of the 180-cm linear treadmill were identified from position resets. Position resets were detected as backward jumps exceeding 20 cm, and position within each lap was independently normalized to span 0–180 cm. A correction was applied to the first lap when recording began partway through the track. Cells were included in place-cell classification if at least five laps and at least 20 APs during movement were available. For classification, spatial tuning curves were calculated from movement periods using 20 equally spaced position bins spanning the 180-cm track. Spatial information^34^ was quantified in bits/spike as follows:

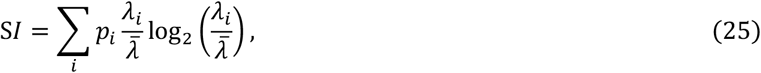

where *p*_*i*_ is the probability of occupying spatial bin i, *λ*_*i*_ is the firing rate in bin i, and 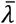 is the mean firing rate across the track. Significance was assessed using 1,000 circularly shifted spike trains. For each shuffle, AP timestamps were shifted relative to behavioural position, the spatial tuning curve was recalculated, and shuffled spatial information was obtained. The empirical one-sided p value was calculated as follows:

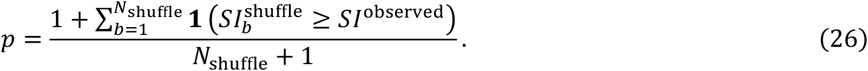

Cells with p < 0.05 were classified as place cells. For visualization of lap-by-lap place fields, the track was divided into 100 bins of 1.8 cm. Occupancy time and spike counts were calculated separately for each lap. Both were smoothed independently using a Gaussian filter with a standard deviation of 9 cm, and the smoothed firing rate was calculated as the ratio of smoothed spike count to smoothed occupancy. Participation of place and non-place cells was compared using the same cell-level binomial framework used for anatomical classes. Differences in membrane-potential and intrinsic-excitability variables were analysed using the corresponding continuous-outcome models. Contributions of Pre depolarization, Ripple depolarization, θ (both θ_mean_ and θ_SD_ together), and R_in_ to the place-cell versus non-place-cell difference in PR were quantified using the same posterior Shapley decomposition described for anatomical cell classes.

#### Rebound analysis

To quantify variation in ripple waveform strength independently of recording-specific differences in LFP scale and waveform shape, ripple-centred LFP traces from −200 to +200 ms around the ripple peak were analysed using principal-component analysis (PCA). Within each recorded cell, the mean ripple waveform was first subtracted from every event. The residual waveforms were then divided by a single standard deviation calculated across all residual waveform values from that cell. The normalized ripple waveforms were pooled across cells and subjected to PCA. The score on the first principal component (PC1) was used as a continuous event-level measure of ripple waveform variation. Because the sign of a principal component is arbitrary, PC1 was oriented such that the mean loading between 0 and 100 ms relative to the ripple peak was positive. For visualization of the corresponding LFP waveforms, events were grouped into five bins according to their PC1 score; all statistical analyses used the continuous PC1 score. Relationships between PC1 and rebound were assessed using hierarchical linear regression with a cell-specific random intercept,

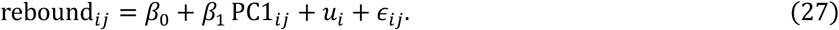

Associations of rebound with ripple amplitude and frequency were analysed using analogous hierarchical regressions. Ripple amplitude and frequency were log-transformed and standardized, and rebound was standardized before model fitting. Ripple participation was related to rebound using a hierarchical Bernoulli regression with both cell-specific intercepts and rebound slopes,

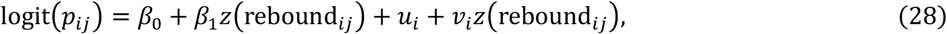

where *u*_*i*_ and *v*_*i*_ represent cell-specific intercept and slope terms, respectively. The relationship between rebound and post-ripple hyperpolarization was assessed using a hierarchical linear regression with a cell-specific random intercept.

To determine the temporal relationship between extracellular and intracellular post-ripple signals, Pearson correlations between LFP and ΔV_m_ were calculated across ripple events separately for each cell and each time point around ripple offset. Correlation coefficients were Fisher z-transformed before averaging across cells and transformed back to Pearson’s r for visualization. Confidence intervals were obtained by resampling cells with replacement 2,000 times. Time periods with a consistent population-level correlation were identified using a cluster-based permutation test. At each time point, Fisher-transformed correlations were tested against zero; contiguous samples with pointwise p<0.05 and the same sign were grouped into clusters, with cluster mass defined as the sum of the absolute t-statistics. A null distribution of the maximum cluster mass was generated from 10,000 random sign-flips at the cell level, and cluster-corrected p-values were calculated from this distribution.

#### Bayesian statistical analysis

Primary statistical analyses were performed in Python using Bambi, PyMC, ArviZ, NumPy, SciPy, pandas, scikit-learn, Pynapple. Individual ripple participation was modelled with Bernoulli likelihoods, whereas cell-level counts of participating and total ripples were modelled using binomial likelihoods. Continuous outcomes on cell-level analyses were modelled predominantly using Student-t likelihoods where robustness to outliers was desirable. Models containing repeated observations from the same neuron included cell-specific random effects as indicated. Bayesian models were sampled using the No-U-Turn Sampler. For the principal cell-level decomposition models, four thousand tuning samples and four thousand posterior samples were drawn with a target acceptance probability of 0.90; target acceptance was increased to 0.95 for selected hierarchical models. Model convergence was evaluated from posterior sampling diagnostics, including 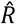, effective sample sizes, and inspection of posterior traces.

Unless otherwise indicated, model parameters and derived quantities are summarized by their posterior median and 95% highest-density interval (HDI). The 95% HDI was used as the Bayesian credible interval and represents the region of highest posterior density containing 95% of the posterior probability. Evidence for an effect differing from the null value was considered when the 95% HDI did not include zero. For quantities expressed as ratios, such as odds ratios, the corresponding null value was one.

## References

1. Buzsáki, G. (2015). Hippocampal sharp wave-ripple: A cognitive biomarker for episodic memory and planning. Hippocampus 25, 1073–1188. 10.1002/hipo.22488.

2. Liu, A.A., Henin, S., Abbaspoor, S., Bragin, A., Buffalo, E.A., Farrell, J.S., Foster, D.J., Frank, L.M., Gedankien, T., Gotman, J., et al. (2022). A consensus statement on detection of hippocampal sharp wave ripples and differentiation from other fast oscillations. Nat. Commun. 13, 6000. 10.1038/s41467-022-33536-x.

3. Foster, D.J., and Wilson, M.A. (2006). Reverse replay of behavioural sequences in hippocampal place cells during the awake state. Nature 440, 680–683. 10.1038/nature04587.

4. O’Neill, J., Senior, T., and Csicsvari, J. (2006). Place-Selective Firing of CA1 Pyramidal Cells during Sharp Wave/Ripple Network Patterns in Exploratory Behavior. Neuron 49, 143–155. 10.1016/j.neuron.2005.10.037.

5. Carr, M.F., Jadhav, S.P., and Frank, L.M. (2011). Hippocampal replay in the awake state: a potential substrate for memory consolidation and retrieval. Nat. Neurosci. 14, 147–153. 10.1038/nn.2732.

6. Dupret, D., O’Neill, J., Pleydell-Bouverie, B., and Csicsvari, J. (2010). The reorganization and reactivation of hippocampal maps predict spatial memory performance. Nat. Neurosci. 13, 995–1002. 10.1038/nn.2599.

7. Gillespie, A.K., Astudillo Maya, D.A., Denovellis, E.L., Liu, D.F., Kastner, D.B., Coulter, M.E., Roumis, D.K., Eden, U.T., and Frank, L.M. (2021). Hippocampal replay reflects specific past experiences rather than a plan for subsequent choice. Neuron 109, 3149–3163.e6. 10.1016/j.neuron.2021.07.029.

8. Ólafsdóttir, H.F., Carpenter, F., and Barry, C. (2017). Task Demands Predict a Dynamic Switch in the Content of Awake Hippocampal Replay. Neuron 96, 925–935.e6. 10.1016/j.neuron.2017.09.035.

9. Pfeiffer, B.E., and Foster, D.J. (2013). Hippocampal place-cell sequences depict future paths to remembered goals. Nature 497, 74–79. 10.1038/nature12112.

10. Van Der Meer, M.A.A., and Bendor, D. (2025). Awake replay: off the clock but on the job. Trends Neurosci. 48, 257–267. 10.1016/j.tins.2025.02.006.

11. Jadhav, S.P., Kemere, C., German, P.W., and Frank, L.M. (2012). Awake Hippocampal Sharp-Wave Ripples Support Spatial Memory. Science 336, 1454–1458. 10.1126/science.1217230.

12. Ylinen, A., Bragin, A., Nadasdy, Z., Jando, G., Szabo, I., Sik, A., and Buzsaki, G. (1995). Sharp wave-associated high-frequency oscillation (200 Hz) in the intact hippocampus: network and intracellular mechanisms. J. Neurosci. 15, 30–46. 10.1523/JNEUROSCI.15-01-00030.1995.

13. English, D.F., Peyrache, A., Stark, E., Roux, L., Vallentin, D., Long, M.A., and Buzsáki, G. (2014). Excitation and Inhibition Compete to Control Spiking during Hippocampal Ripples: Intracellular Study in Behaving Mice. J. Neurosci. 34, 16509–16517. 10.1523/JNEUROSCI.2600-14.2014.

14. Hulse, B.K., Moreaux, L.C., Lubenov, E.V., and Siapas, A.G. (2016). Membrane Potential Dynamics of CA1 Pyramidal Neurons during Hippocampal Ripples in Awake Mice. Neuron 89, 800–813. 10.1016/j.neuron.2016.01.014.

15. Noguchi, A., Huszár, R., Morikawa, S., Buzsáki, G., and Ikegaya, Y. (2022). Inhibition allocates spikes during hippocampal ripples. Nat. Commun. 13, 1280. 10.1038/s41467-022-28890-9.

16. Valero, M., Cid, E., Averkin, R.G., Aguilar, J., Sanchez-Aguilera, A., Viney, T.J., Gomez-Dominguez, D., Bellistri, E., and de la Prida, L.M. (2015). Determinants of different deep and superficial CA1 pyramidal cell dynamics during sharp-wave ripples. Nat. Neurosci. 18, 1281–1290. 10.1038/nn.4074.

17. Baimbridge, K.G., Peet, M.J., McLennan, H., and Church, J. (1991). Bursting response to current-evoked depolarization in rat ca1 pyramidal neurons is correlated with lucifer yellow dye coupling but not with the presence of calbindin-D28k. Synapse 7, 269–277. 10.1002/syn.890070404.

18. Aleman-Zapata, A., Capitan, M.M., Samanta, A., Özsezer, P., Agarwal, K., Adam, T., Rayan, A., and Genzel, L. (2025). Differential contributions of CA3 and entorhinal cortex inputs to ripple patterns in the hippocampus. iScience 28, 111782. 10.1016/j.isci.2025.111782.

19. Castelli, M., Lopes-dos-Santos, V., Gava, G.P., Lambiotte, R., and Dupret, D. (2025). Hippocampal ripple diversity organizes neuronal reactivation dynamics in the offline brain. Neuron 113, 4245–4262.e17. 10.1016/j.neuron.2025.09.012.

20. Ramirez-Villegas, J.F., Logothetis, N.K., and Besserve, M. (2015). Diversity of sharp-wave–ripple LFP signatures reveals differentiated brain-wide dynamical events. Proc. Natl. Acad. Sci. 112, E6379–E6387. 10.1073/pnas.1518257112.

21. Sebastian, E.R., Quintanilla, J.P., Sánchez-Aguilera, A., Esparza, J., Cid, E., and De La Prida, L.M. (2023). Topological analysis of sharp-wave ripple waveforms reveals input mechanisms behind feature variations. Nat. Neurosci. 26, 2171–2181. 10.1038/s41593-023-01471-9.

22. Paleologos, N., Vöröslakos, M., Gonzalez, J., Maslarova, A., Aykan, D., Liu, A.A., and Buzsáki, G. (2025). Electroanatomy of hippocampal activity patterns: theta, gamma waves, sharp wave-ripples, and dentate spikes. Front. Behav. Neurosci. 19, 1685846. 10.3389/fnbeh.2025.1685846.

23. Mizuseki, K., Diba, K., Pastalkova, E., and Buzsáki, G. (2011). Hippocampal CA1 pyramidal cells form functionally distinct sublayers. Nat. Neurosci. 14, 1174–1181. 10.1038/nn.2894.

24. Abbaspoor, S., and Hoffman, K.L. (2024). Circuit dynamics of superficial and deep CA1 pyramidal cells and inhibitory cells in freely moving macaques. Cell Rep. 43, 114519. 10.1016/j.celrep.2024.114519.

25. O’Keefe, J., and Dostrovsky, J. (1971). The hippocampus as a spatial map. Preliminary evidence from unit activity in the freely-moving rat. Brain Res. 34, 171–175. 10.1016/0006-8993(71)90358-1.

26. Henze, D.A., and Buzsáki, G. (2001). Action potential threshold of hippocampal pyramidal cells in vivo is increased by recent spiking activity. Neuroscience 105, 121–130. 10.1016/S0306-4522(01)00167-1.

27. Platkiewicz, J., and Brette, R. (2011). Impact of Fast Sodium Channel Inactivation on Spike Threshold Dynamics and Synaptic Integration. PLOS Comput. Biol. 7, e1001129. 10.1371/journal.pcbi.1001129.

28. Rangel Guerrero, D.K., Balueva, K., Barayeu, U., Baracskay, P., Gridchyn, I., Nardin, M., Roth, C.N., Wulff, P., and Csicsvari, J. (2024). Hippocampal cholecystokinin-expressing interneurons regulate temporal coding and contextual learning. Neuron 112, 2045–2061.e10. 10.1016/j.neuron.2024.03.019.

29. Fuentealba, P., Begum, R., Capogna, M., Jinno, S., Márton, L.F., Csicsvari, J., Thomson, A., Somogyi, P., and Klausberger, T. (2008). Ivy Cells: A Population of Nitric-Oxide-Producing, Slow-Spiking GABAergic Neurons and Their Involvement in Hippocampal Network Activity. Neuron 58, 295. 10.1016/j.neuron.2008.04.004.

30. Bittner, K.C., Grienberger, C., Vaidya, S.P., Milstein, A.D., Macklin, J.J., Suh, J., Tonegawa, S., and Magee, J.C. (2015). Conjunctive input processing drives feature selectivity in hippocampal CA1 neurons. Nat. Neurosci. 18, 1133–1142. 10.1038/nn.4062.

31. Epsztein, J., Brecht, M., and Lee, A.K. (2011). Intracellular Determinants of Hippocampal CA1 Place and Silent Cell Activity in a Novel Environment. Neuron 70, 109–120. 10.1016/j.neuron.2011.03.006.

32. Zhang, X., Schlögl, A., and Jonas, P. (2020). Selective Routing of Spatial Information Flow from Input to Output in Hippocampal Granule Cells. Neuron 107, 1212–1225.e7. 10.1016/j.neuron.2020.07.006.

33. Mensi, S., Naud, R., Pozzorini, C., Avermann, M., Petersen, C.C.H., and Gerstner, W. (2012). Parameter extraction and classification of three cortical neuron types reveals two distinct adaptation mechanisms. J. Neurophysiol. 107, 1756–1775. 10.1152/jn.00408.2011.

34. Skaggs, W., McNaughton, B., and Gothard, K. (1992). An information-theoretic approach to deciphering the hippocampal code. In Advances in neural information processing systems, S. Hanson, J. Cowan, and C. Giles, eds. (Morgan-Kaufmann).

