## Supplemental Figures for "Cellular Determinants of CA1 Pyramidal Cell Recruitment during Sharp-Wave Ripples"

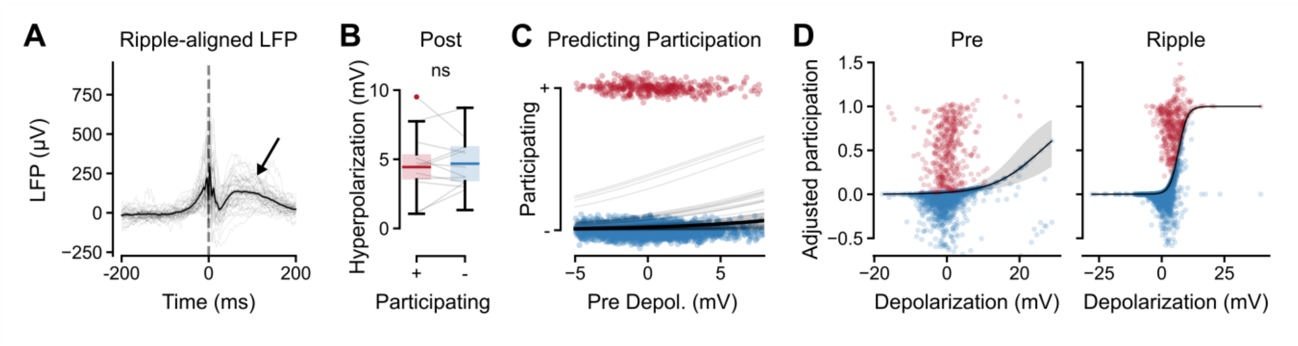


**Supplementary Figure 1: Additional analyses on membrane-potential dynamics underlying CA1 pyramidal-cell participation during hippocampal ripples**

(**A**) LFP traces aligned to the ripple peak. Thin lines indicate individual cell averages, and the black line indicates the population average. Arrow highlights post-ripple rebound.

(**B**) Comparison of post-ripple hyperpolarization between participating (+) and non-participating (−) events across cells.

(**C**) Relationship between pre-depolarization and the probability of participation. Points show individual ripple events and curves show cell-specific participation functions estimated using hierarchical logistic regression.

**(D)** Relationship between pre- and ripple-depolarization and participation probability after adjusting for the other depolarization measure and cell-specific random effects. Points represent individual ripple events plotted as partial residuals on the probability scale, and curves show the fitted population-level relationship.

Statistical evidence is denoted by asterisks according to the highest credible level whose highest density interval excludes the null (*, 95%; **, 99%; ***, 99.9%; ns, otherwise).

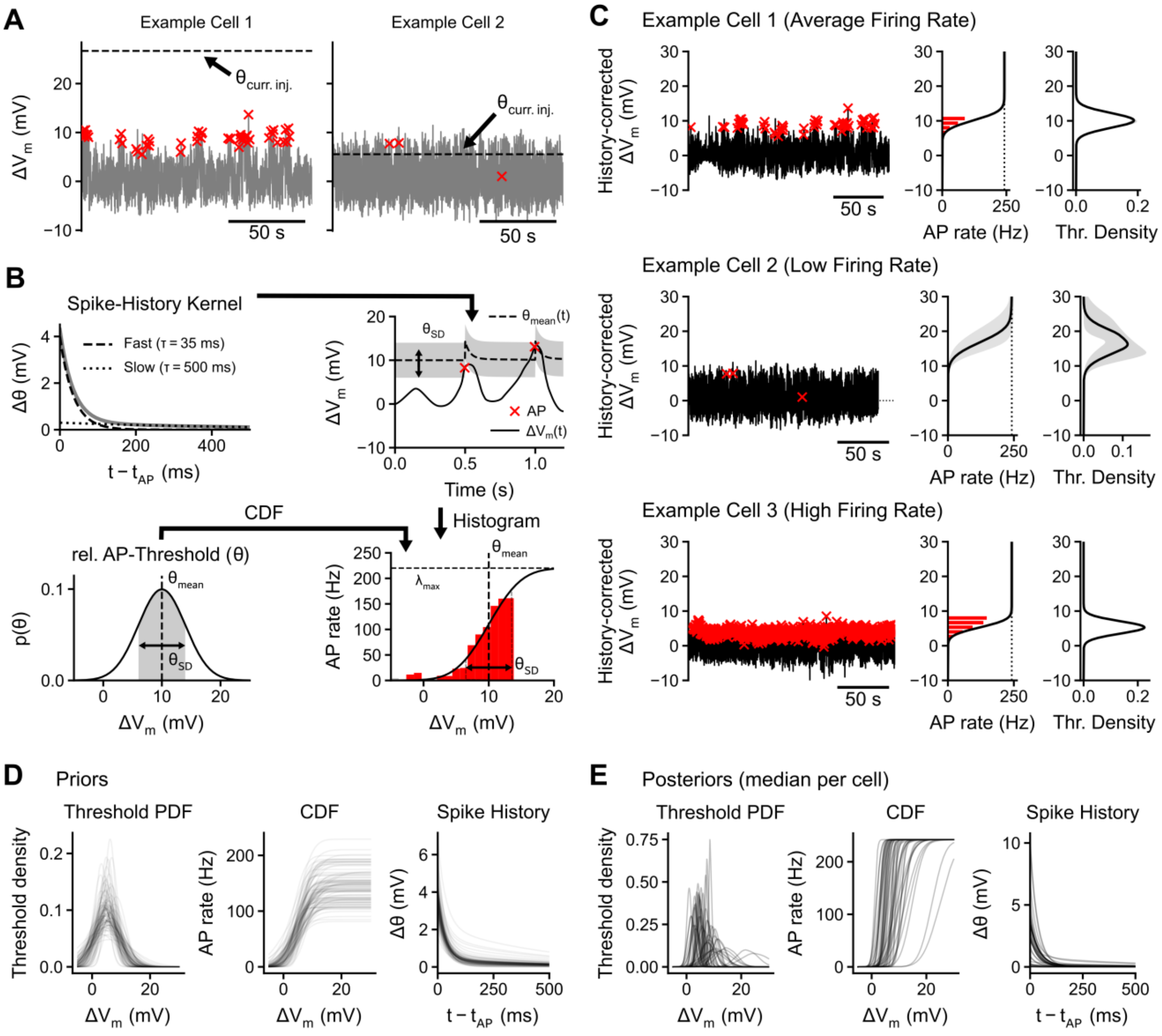


**Supplementary Figure 2**: **Probabilistic threshold model of AP generation.**

**(A)** $\Delta V_m$ traces of two example neurons where the AP-Threshold estimated from current injections ($\theta_{curr.inj.}$) did not match the data during spontaneous activity. AP-onset is marked by red crosses.

**(B)** Schematic of the model. Relative action-potential threshold, θ, was modelled as a Gaussian distribution with cell-specific mean ($\theta_{mean}$) and standard deviation ($\theta_{SD}$). The corresponding cumulative distribution function (CDF) maps history-corrected membrane-potential depolarization, $\Delta V_m$, onto instantaneous AP rate, which saturates at the maximum firing rate ($\lambda_{max}$). Spike history was incorporated using fast and slow exponentially decaying kernels ($\tau_{fast}$= 35 ms; $\tau_{fast}$= 500 ms), transiently shifting the effective threshold following an AP. Example trace illustrate the resulting time-varying threshold and AP generation.

**(C)** Model fits for three example cells spanning different firing-rate regimes. Left, history-corrected $\Delta V_m$ traces with detected APs indicated in red. Middle, estimated relationship between $\Delta V_m$ and AP rate; red bars show the empirical binned AP rate, and the black curve shows the fitted CDF-based rate function. Dotted vertical lines indicate $\lambda_{max}$. Right, fitted threshold distributions for each cell; gray shading indicates posterior uncertainty. Example Cells 1&2 correspond to the cells shown in (A).

**(D)** Prior distributions for the main model components. Left, prior predictive distributions of the Gaussian threshold density. Middle, corresponding CDF-based AP-rate functions. Right, prior distributions of the spike-history kernel. Thin lines indicate individual prior draws.

**(D)** Posterior estimates of the same model components, summarized at the cell level. Left, posterior median threshold-density functions for individual cells. Middle, posterior median CDF-based AP-rate functions. Right, posterior median spike-history kernels. Each line represents one cell, illustrating substantial cell-to-cell variation in threshold position and width, AP-rate transformation, and spike-history effects.

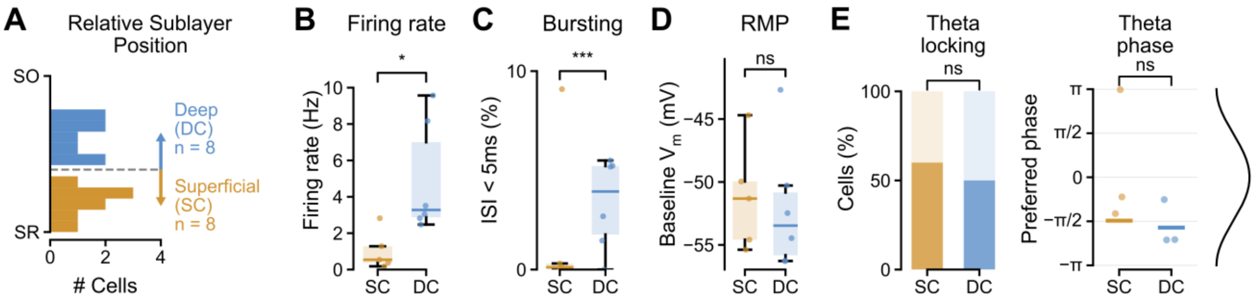


**Supplementary Figure 3: Functional properties of superficial and deep CA1 pyramidal cells**

**(A)** Histogram of the relative sublayer position of superficial (SC) and deep cells (DC). Cutoff border (0.4) is indicated using a dashed line.

**(B)** Firing rate differences between superficial and deep cells. Difference was tested using negative binomial regression.

**(C)** AP-burst likelihood differences between superficial and deep cells. AP bursts were defined as an inter-spike interval (ISI) less than 5 ms. Difference was tested using binomial regression.

**(D)** Resting membrane potential (RMP) differences between superficial and deep cells, measured as the baseline $V_m$. Difference was tested using Student-t regression.

**(E)** Theta phase-locking properties of superficial and deep cells. Left: Proportion of superficial and deep cells with significant theta phase locking (mean vector length > 0.1). Difference between groups was tested using a Fisher's exact test. Right: Preferred phase of cells with significant phase-locking. Difference in preferred phase was tested using a permutation test.

Statistical evidence is denoted by asterisks according to the highest credible level whose highest density interval excludes the null (*, 95%; **, 99%; ***, 99.9%; ns, otherwise).

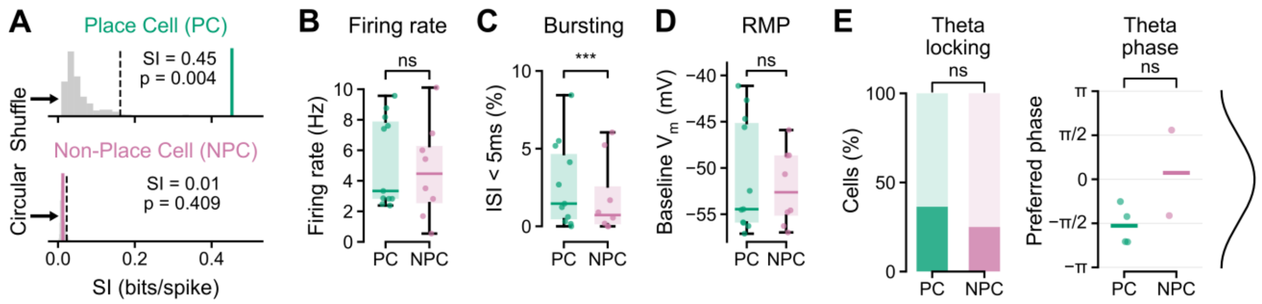


**Supplementary Figure 4: Functional properties of place and non-place cells**

**(A)** Definition of place cells. Spatial information (SI) was calculated from the AP-rate map and compared to a null distribution (grey histogram) estimated using circular shuffling of spike times. If the observed SI (solid line) exceeded the 95$^{th}$ percentile of said distribution (dashed line) the cell was classified as a place cell (PC; top) or a non-place cell otherwise (NPC; bottom).

**(B)** Firing rate differences between place and non-place cells. Difference was tested using negative binomial regression.

**(C)** AP-burst likelihood differences between place and non-place cells. AP bursts were defined as an inter-spike interval (ISI) less than 5 ms. Difference was tested using binomial regression.

**(D)** Resting membrane potential (RMP) differences between place and non-place cells, measured as the baseline $V_m$. Difference was tested using Student-t regression.

**(E)** Theta phase-locking properties of place and non-place cells. Left: Proportion of place and non-place cells with significant theta phase locking (mean vector length > 0.1). Difference between groups was tested using a Fisher's exact test. Right: Preferred phase of cells with significant phase-locking. Difference in preferred phase was tested using a permutation test.

Statistical evidence is denoted by asterisks according to the highest credible level whose highest density interval excludes the null (*, 95%; **, 99%; ***, 99.9%; ns, otherwise).
